# Loss of Sall3 eliminates presynaptic inhibition of sensory motor circuits and impairs adaptive motor behavior

**DOI:** 10.64898/2026.08.25.746905

**Authors:** Jennifer L. Shadrach, Amr A. Mahrous, Robert Palovics, Zinnia Saha, Richard H. Roth, Atharv Panditrao, Vanessa W.Y. Kan, Mark A. Gradwell, Victoria E. Abraira, Irene L. Llorente, Jun B. Ding, Tony Wyss-Coray, David J. Bennett, CJ Heckman, Julia A. Kaltschmidt

## Abstract

Spinal presynaptic inhibitory interneurons are thought to regulate proprioceptive sensory feedback to shape motor output, however, their specific contribution to motor behavior has been difficult to assess, partially due to the lack of a specific genetic handle. Here, we identify Sall3 as the transcription factor required for the establishment and maintenance of GABApre axo-axonic synapses on proprioceptive Ia afferent terminals. Loss of Sall3 in mice selectively eliminates GABApre boutons on Ia afferent terminals, resulting in altered sensory-evoked motor responses and impaired skilled locomotor behaviors. Together, these findings establish Sall3 as a key regulator of GABApre circuit development and provide a genetic framework for understanding how presynaptic inhibition shapes sensorimotor integration.

## Introduction

Coordinated movement depends on continuous sensory feedback to inform spinal motor circuits about body position and the external environment. A central component of this control is the monosynaptic stretch reflex, in which group Ia proprioceptive afferents provide direct excitatory input to motor neurons.^1–3^ The gain of this pathway is regulated by a specialized population of presynaptic GABAergic (GABApre) interneurons that form axoaxonic boutons onto Ia afferent terminals.^4–6^ By modulating Ia afferent glutamate release, GABApre interneurons thereby regulate excitatory drive onto motor neurons and shape reflex output.^7–9^ Presynaptic inhibition is thought to enable context-dependent control of sensory feedback during movement while preventing excessive reflex activation.^10–12^ However, the specific contribution of presynaptic inhibition to motor behavior has been difficult to assess.

GABApre interneurons belong to a broader set of spinal presynaptic inhibitory interneurons generated from Ptf1a-defined dorsal progenitor domains.^13–17^ The identification of selective molecular markers has enabled detailed studies of the development, connectivity, and function of several members of this broader class. For example, parvalbumin- and calretinin-expressing interneurons mediate presynaptic inhibition of cutaneous afferents,^18,19^ and Rorb-expressing interneurons that gate proprioceptive input in the deeper dorsal horn.^20^ In contrast, GABApre interneurons lack a comparably selective molecular identifier. Consequently, studies of GABApre circuitry have instead relied on broader features associated with GABApre identity, including Ptf1a lineage tracing^6,21–23^ and expression of the GABA-synthetic enzyme glutamic acid decarboxylase (GAD)65, encoded by *Gad2*, which is highly enriched in GABApre boutons on Ia terminals.^5,6,12^ Moreover, although several genes have been implicated in GABApre synapse formation, loss-of-function studies produce only partial reductions in presynaptic boutons on Ia afferent terminals,^21,23^ suggesting that additional factors contribute to the establishment of these synapses.

The inability to selectively and completely manipulate GABApre interneurons has also complicated efforts to resolve how presynaptic inhibition regulates sensory-motor integration and contributes to locomotor control. Classical models attribute presynaptic inhibition to primary afferent depolarization (PAD), in which activation of GABA_A_ receptors at Ia afferent terminals is thought to depolarize the terminal and suppress neurotransmitter release.^7,24^ However, recent evidence indicates that GABA_A_ receptors are largely excluded from Ia afferent terminals and instead localize to nodes of Ranvier along sensory axons, where their activation can facilitate sensory transmission.^8,9^ In contrast, GABA_B_ receptors at Ia afferent terminals are thought to mediate suppression of neurotransmitter release.^9,25^ This suggests that PAD and presynaptic inhibition may be mediated by different neuronal populations that operate at distinct sites along the sensory axon. Defining the molecular basis of presynaptic inhibition will therefore be essential for uncovering the circuit logic by which sensory feedback shapes motor output and supports stable yet adaptable locomotion.

Here, we identify the transcription factor *Sal-like protein 3* (*Sall3*) as a Ptf1a-dependent marker enriched in dI4/LA interneurons. Using Sall3 loss-of-function mice, we show for the first time the elimination of GABApre axoaxonic boutons on Ia afferent terminals in the ventral spinal cord. Despite this loss of presynaptic terminals, PAD is largely intact, providing genetic evidence that PAD arises from sites outside Ia afferent terminals. Sall3 mutants further exhibit altered sensory-evoked motor responses and impairments skilled locomotor and swimming behaviors. Together, these findings establish Sall3 as a critical regulator of GABApre circuit development and provide new insight into the relationship between presynaptic inhibition, sensory processing, and adaptive motor control.

## Results

### Developmental single-cell analysis identifies Sall3 as a marker of dI4/LA interneurons

To identify molecular markers of GABApre interneurons, we analyzed a published single-cell RNA sequencing dataset of developing mouse spinal cord spanning embryonic days E9.5-E13.5,^26^ focusing on dI4/LA interneurons that give rise to GABApre interneurons.^6,23^ We first clustered and annotated broad cell classes (see Methods; Figures S1A-G), followed by interneuron reclustering. This analysis resolved 19 transcriptionally distinct populations that mapped to established cardinal interneuron classes based on canonical transcription factor expression (Figures 1A-D).^26–29^ Among these populations, inhibitory dI4/LA and excitatory dI5/LB interneurons were the most abundant (Figure 1C), consistent with the extended period of dorsal neurogenesis that generates late-born dILA and dILB neurons.^30–32^ The relative abundance of dI4/LA interneurons enabled high-resolution characterization of molecular heterogeneity within this population (Figures SI-L).

**Figure 1.**
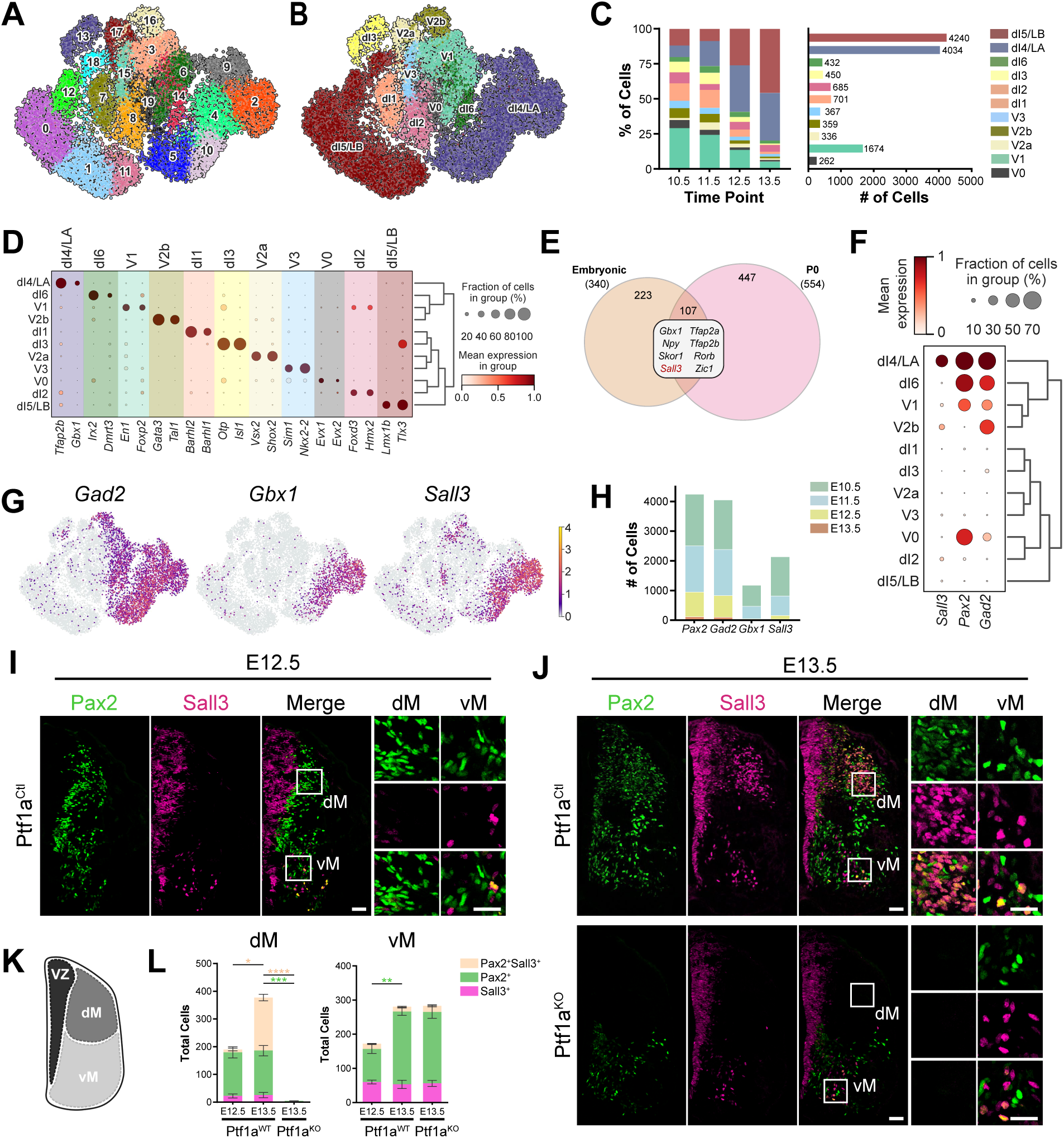
Sall3 expression in dI4/LA interneurons requires Ptf1a. **(A-B)** UMAP projections of all interneuron populations from the Delilie et al. embryonic spinal cord dataset^26^ showing cluster assignments (A) and annotated cell type identities (B). **(C)** Proportion of interneurons at each time point (left) and total cell number (right) for each annotated interneuron cluster. **(D)** Dot plot showing expression of canonical transcription factors across annotated spinal interneuron lineages. **(E)** Comparison of genes enriched in embryonic dI4/LA interneurons with those reported in a postnatal spinal cord dataset from Osseward et al.^33^ revealed 107 shared genes and highlighted *Sall3* as a candidate dI4/LA marker. **(F)** Dot plot showing *Sall3* expression is selectively enriched in *Pax2*^+^*Gad2*^+^ dI4/LA interneurons. **(G-H)** *Gad2*, *Gbx1*, and *Sall3* expression in UMAP space (G). *Gbx1* and *Sall3* show a similar developmental enrichment at later embryonic stages (H). **(I-J)** Immunostaining for Pax2 (green) and Sall3 (magenta) in representative E12.5 Ptf1a^Ctl^ (I), E13.5 Ptf1a^Ctl^ (J, top), and E13.5 Ptf1a^KO^ (J, bottom) embryonic spinal cords. Insets show higher-magnification views of regions in the dM and vM. **(K)** Schematic of the embryonic spinal cord. **(L)** Quantification of Pax2 and Sall3 co-expression in the dM (left) and vM (right) of Ptf1a^Ctl^ and Ptf1a^KO^ embryos. All scale bars, 50 µm. Graphs show mean ± SEM. Statistical significance is indicated as * *p* < 0.05, ** *p* < 0.01, *** *p* < 0.001, **** *p* < 0.0001. For panel L, asterisks are color-matched to the quantified cell population. Exact *p* values, sample sizes (N), means, and standard deviations are provided in Table S1. Abbreviations: dI, dorsal interneuron; v, ventral interneuron; VZ, ventricular zone; dM, dorsal mantle zone; vM, ventral mantle zone.

To identify genes enriched in dI4/LA interneurons, we performed differential gene expression analysis relative to V1 inhibitory interneurons, leveraging their shared GABAergic identity to exclude broadly expressed inhibitory genes (Figure S1H). We then intersected this embryonic dI4/LA-enriched gene set with genes enriched in postnatal dI4/LA interneurons,^33^ yielding 107 shared genes (Figure 1E). After excluding transcripts with broader interneuron expression patterns (such as *Zic1* and *Skor1*^32,34^) as well as established dI4/LA markers (including *Tfap2b*, *Npy*, *Gbx1*, and *Rorb*^13,16,17,20,35–37)^, *Sall3* emerged as a previously uncharacterized gene enriched in dI4/LA interneurons (Figure 1F).

Because GABApre interneurons arise from late-born dILA interneurons,^23^ we first asked whether Sall3 expression was consistent with this developmental origin. *Sall3* expression closely aligned with the dILA marker *Gbx1*^36^ in UMAP space (Figure 1G). In addition, *Sall3* and *Gbx1* were preferentially expressed at later embryonic stages (E12.5-E13.5; Figure 1H), whereas pan-GABAergic markers *Pax2* and *Gad2* were broadly expressed across dI4/LA populations throughout embryonic development.

During spinal cord development, neural progenitors in the ventricular zone generate distinct neuronal lineages that migrate into the dorsal and ventral mantle zones, where they continue to differentiate into mature interneuron subtypes (Figure 1K).^29^ To determine if Sall3 protein expression mirrors the transcriptomic profile predicted by the single-cell data, we examined Sall3 and Pax2 co-expression at embryonic stages. At E12.5, Sall3 was sparsely expressed in 6% of Pax2^+^ interneurons in the dorsal mantle zone (Figures 1I,L). By E13.5, however, Sall3 expression had increased markedly, and 54% of dorsal Pax2^+^ interneurons were Sall3^+^ (Figures 1J,L). This delayed accumulation of Sall3 protein mirrors its late embryonic transcriptomic expression and further supports an association with late-born dILA population.

We next asked whether Sall3 expression depends on *Ptf1a*, a transcription factor required for specification of dorsal dI4/LA interneurons.^14^ Ptf1a^KO^ embryos failed to generate dorsal Pax2^+^ interneurons, and consistent with a dI4/LA origin, Sall3 expression was correspondingly absent from the dorsal mantle zone (Figures 1J-L). Notably, more than 90% of Pax2^+^Sall3^+^ interneurons were located within the dorsal mantle zone at E13.5, whereas only 6% of ventral Pax2^+^ interneurons expressed Sall3 (Figure 1L). The small ventral population was retained in Ptf1a^KO^ embryos, indicating that the predominant dorsal Sall3^+^ population arises from a Ptf1a-dependent lineage.

Finally, we asked whether postnatal Sall3 expression matched the known anatomical distribution of GABApre interneurons. After birth, the spinal cord retains a clear dorsoventral organization, with the superficial dorsal and deep dorsal horn (DH, DDH) mediating sensory integration and the ventral horn (VH) controlling motor output (Figure 2A). Previous studies localized GABApre interneurons to the medial DDH (laminae V-VI), adjacent to the central canal, at postnatal stages.^5,6,38^ Consistent with this distribution, Sall3 expression was retained throughout postnatal development in dorsal and deep dorsal Pax2^+^ interneuron populations (DH: 80%; DDH: 60% at P0, 40% at P25), whereas in the VH Sall3 was limited to a small fraction of Pax2^+^ interneurons and was nearly absent by P25 (Figures 2B,C; Figure S2E). Spatial mapping within dorsal domains further showed that although Sall3^+^Pax2^+^ interneurons were broadly distributed across the DH, they were preferentially localized to the medial DDH (Figures 2D,E; Figures S2H,I). Together, these findings support *Sall3* as a marker of a dI4/LA subpopulation that includes the presumptive GABApre lineage, motivating functional analysis of *Sall3* in GABAergic interneurons.

**Figure 2.**
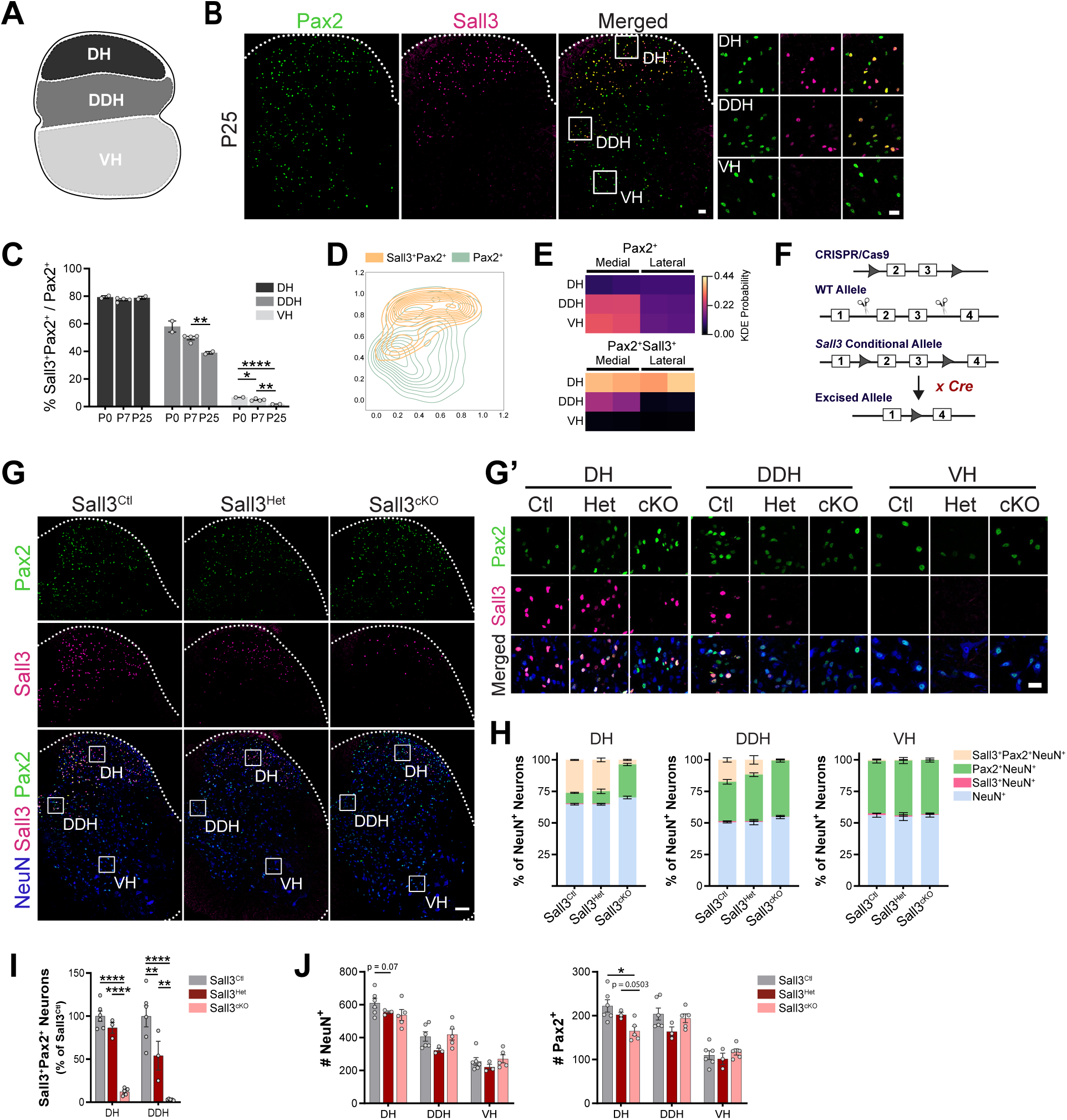
Conditional deletion of Sall3 in GABAergic interneurons. **(A)** Schematic of the postnatal spinal cord. **(B)** Immunostaining for Pax2 (green) and Sall3 (magenta) in a representative P25 spinal cord hemisection. Insets show higher-magnification views of regions in the DH, DDH, and VH. Representative images for P0 and P7 are shown in Figure S2E. **(C)** Quantification of Sall3 and Pax2 co-expression across postnatal development. The percentage of Sall3^+^Pax2^+^ cells relative to the total Pax2^+^ population is shown for DH (black), DDH (dark gray), and VH (light gray) at P0, P7, and P25. **(D-E)** Contour plots show the relative density of Sall3^+^Pax2^+^ (orange) and Pax2^+^ (green) interneurons based on normalized coordinates (D). Heatmaps show kernel density estimation (KDE) probability within the DH, DDH, and VH for each population (E). **(F)** Generation of the *Sall3* conditional allele and strategy for selective deletion. Exons 2 and 3 were flanked by loxP sites using CRISPR/Cas9-mediated genome editing. Crossing *Sall3^fx/fx^* mice with a *Gad2^Cre^* driver line results in deletion of *Sall3* in postmitotic GABAergic interneurons. **(G)** Immunostaining for Pax2 (green), Sall3 (magenta), and NeuN (blue) in representative P25 spinal cord hemisections from Sall3^Ctl^, Sall3^Het^, and Sall3^cKO^ mice. Insets (G’) show higher-magnification views of regions in the DH, DDH, and VH. **(H)** Stacked bar plots showing the proportional contribution of neuronal subpopulations relative to the total NeuN^+^ neurons in DH (left), DDH (middle), and VH (right). Cell classes include: Sall3^+^Pax2^+^NeuN^+^ (peach), Pax2^+^NeuN^+^ (green), Sall3^+^NeuN^+^ (magenta), NeuN^+^ (blue). **(I)** Quantification of Sall3^+^Pax2^+^ neurons in the DH and DDH, expressed as the percentage of the corresponding Sall3^Ctl^ value. **(J)** Quantification of NeuN^+^ neurons (left) and Pax2^+^ interneurons (right) per section across the DH, DDH, and VH for all three genotypes. Scale bars: (B) 50 µm (overview), 25 µm (insets); (G) 100 µm (overview), 25 µm (G’ insets). Graphs show mean ± SEM, with dots representing individual mice. Statistical significance is indicated as ** *p* < 0.01, *** *p* < 0.001, **** *p* < 0.0001. Exact *p* values, sample sizes (N), means, and standard deviations are provided in Table S1. Abbreviations: DH, dorsal horn; DDH, deep dorsal horn; VH, ventral horn.

### Conditional deletion of Sall3 in GABAergic interneurons

To investigate the requirement for *Sall3* in spinal inhibitory interneurons, we generated a conditional *Sall3* mouse line (Figure 2F). Crossing this line with a *Gad2^Cre^* driver restricted deletion to postmitotic GABAergic interneurons, thereby avoiding effects on Sall3-expressing neural progenitors and astrocytes (Figures S2A-D,F-G). This strategy generated Sall3^cKO^ (*Gad2^Cre/+^*;*Sall3^fx/fx^*), Sall3^Het^ (*Gad2^Cre/+^*;*Sall3^fx/+^*), and Sall3^Ctl^ (*Gad2^+/+^*;*Sall3^fx/fx^* or *Gad2^+/+^*;*Sall3^fx/+^*) littermates. Sall3^cKO^ mice were recovered at expected Mendelian ratios and exhibited only a mild reduction in body weight (Figure S3A).

We first performed immunostaining to confirm Sall3 deletion in Sall3^cKO^ mice and assess for gross changes in neuronal numbers (Figure 2G, insets 2G’). In the DDH, Sall3 expression was nearly absent from Pax2^+^ interneurons in Sall3^cKO^ mice and reduced in Sall3^Het^ animals, consistent with gene dosage sensitivity (Figures 2H,I). In the DH, a small number of Pax2^+^ interneurons retained Sall3 expression, likely arising from GABAergic or glycinergic interneurons outside the *Gad2* lineage.^39^ In the absence of Sall3, NeuN^+^ neurons and Pax2^+^ interneurons were largely maintained, with only a small, region-specific reduction in DH Pax2^+^ interneurons (Figure 2J).

The sustained enrichment of Sall3 in dorsal inhibitory neurons, together with previous reports linking Ptf1a-lineage interneurons to pathological itch,^40,41^ prompted us to evaluate somatosensory behaviors in Sall3^cKO^ mice. Approximately 50% of Sall3^cKO^ mice developed severe skin lesions, typically at the base of the tail and hindlimb region (Figure S3B). Sall3^cKO^ mice also exhibited increased spontaneous scratching (Figure S3C) and significantly enhanced responses to chemically induced itch (Figures S3D,E). We also examined nociception and found that capsaicin-evoked behaviors and phase I formalin responses were normal, indicating intact acute pain processing (Figures S3F-H). However, phase II formalin response, which reflect persistent nociceptive processing following inflammatory injury, were significantly reduced in Sall3^cKO^ mice (Figure S3H). No additional differences were detected across other somatosensory modalities, including mechanical (von Frey) and thermal (hot plate and dry ice) sensation (Figures S3I,J).

### Formation of GABApre boutons on group Ia proprioceptive afferents requires Sall3

We next focused on presynaptic inhibition of Ia afferent processing. To determine whether loss of Sall3 disrupts the formation of this inhibitory microcircuit, we examined association of GABAergic boutons with sensory afferent terminals in Sall3^Ctl^, Sall3^Het^, and Sall3^cKO^ mice. Sensory afferents were identified by vesicular glutamate transporter (vGlut)1 expression, whereas GAD65 (G65), which is selectively enriched in GABApre boutons, was used to visualize presynaptic inhibitory contacts (Figures 3A-C).^6,12^ We reconstructed G65 puncta contacting vGlut1^+^ terminals (G65^on^; Figure 3D) and quantified their abundance and morphological properties within the DH, DDH, and two VH regions (v1 and v2; see Figures 3A-C insets), which span distinct dorsoventral positions and motor pool territories.^42^

**Figure 3.**
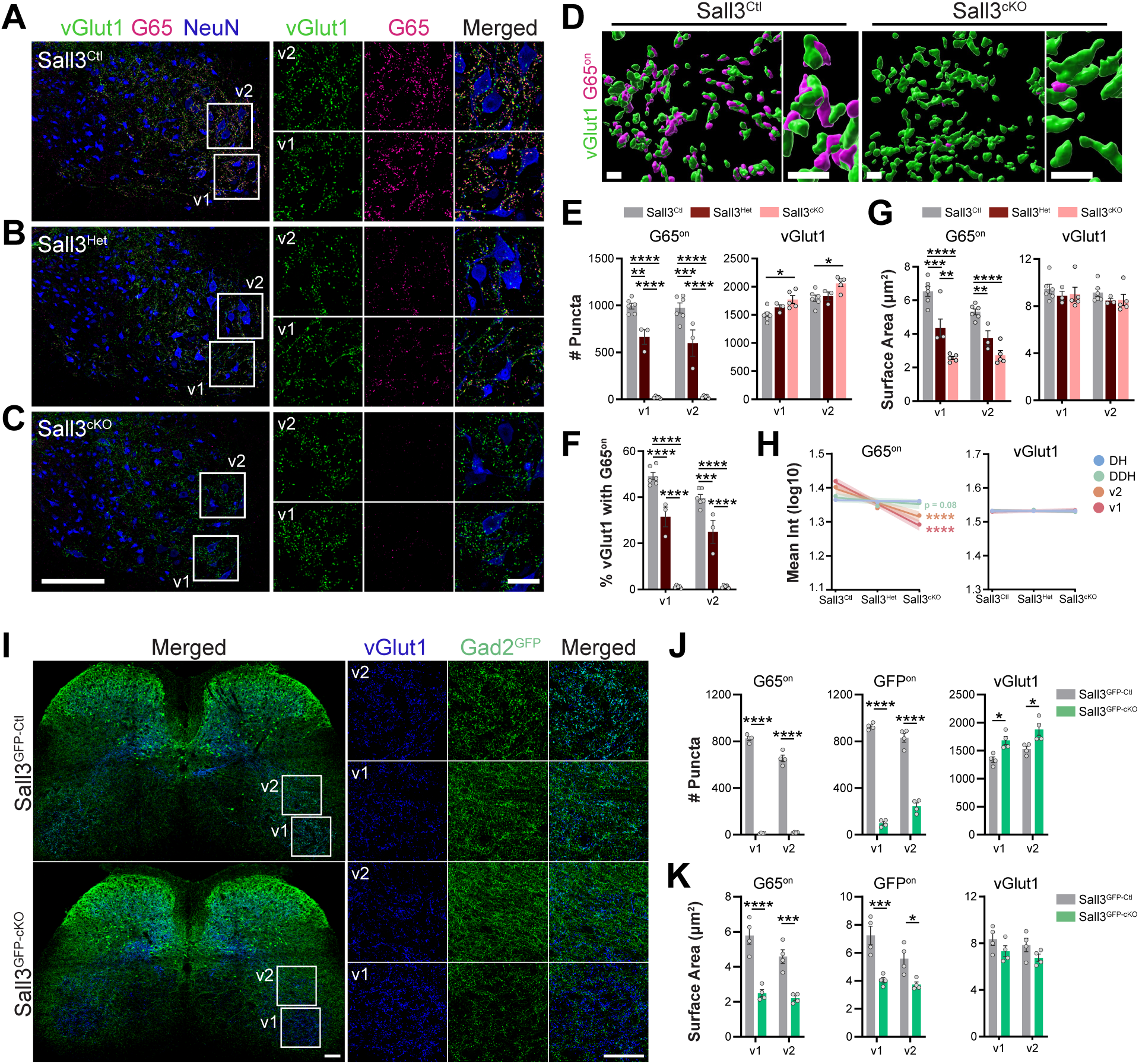
Formation of GABApre boutons on group Ia proprioceptive afferents requires Sall3. **(A-C)** Immunostaining for vGlut1 (green, Ia afferent terminals), G65 (GAD65; magenta, GABApre boutons), and NeuN (blue, neurons) in the ventral horn of representative Sall3^Ctl^ (A), Sall3^Het^ (B), and Sall3^cKO^ (C) spinal cord sections. Insets show higher-magnification views of two ventral horn regions corresponding to v1 and v2. **(D)** Synapse reconstructions showing vGlut1 surfaces (green) and G65 surfaces directly contacting vGlut1 surfaces (G65^on^, magenta) in Sall3^Ctl^ and Sall3^cKO^ mice. **(E)** Number of reconstructed G65^on^ and vGlut1 puncta per section in the v1 and v2 regions across all genotypes. **(F)** Percentage of vGlut1 surfaces contacting at least one associated G65^on^ surface. **(G)** Mean surface area of reconstructed G65^on^ and vGlut1 puncta per section in the v1 and v2 regions across all genotypes. **(H)** Mean fluorescence intensity (Int) of reconstructed surfaces plotted as a function of gene dose with fitted trends. **(I)** Immunostaining for vGlut1 (blue, Ia afferent terminals) and Gad2^GFP^ (green, projections from *Gad2*-expressing neurons) in representative P25 spinal cord sections from Sall3^GFP-Ctl^ and Sall3^GFP-cKO^ mice. Insets show higher-magnification views of v1 and v2 regions. **(J-K)** Quantification of synapse reconstructions from Gad2^GFP^ experiments. Number of puncta per section (J) and mean surface area (K) are shown for G65^on^, GFP^on^, and vGlut1. Scale bars: (A-C) 100 µm (overview), 50 µm (insets); (D) 3 µm (overview and inset); (I) 100 µm (overview and inset). Graphs show mean ± SEM, with dots representing individual mice. Statistical significance is indicated as * *p* < 0.05, ** *p* < 0.01, *** *p* < 0.001, **** *p* < 0.0001. Exact *p* values, sample sizes (N), means, and standard deviations are provided in Table S1.

Loss of Sall3 resulted in a profound depletion of GABApre boutons throughout the spinal cord. G65^on^ puncta were reduced in the DH (58.0 ± 8.8%) and DDH (77.1 ± 6.9%) (Figures S4A-F), but the effect was most severe in the VH, where GABApre boutons were nearly eliminated. In the VH, G65^on^ puncta number decreased by 98.0 ± 1.2% in v1 and 97.3 ± 1.4% in v2 (Figure 3E), and the fraction of vGlut1^+^ terminals receiving at least one GABApre contact fell from ∼40-50% in controls to ∼1% in Sall3^cKO^ mice (Figure 3F). The few remaining G65^on^ puncta in Sall3^cKO^ mice had reduced surface area (∼50-60% decrease) and fluorescence intensity (Figures 3G,H), consistent with a marked reduction in G65 enrichment and diminished GABAergic signaling capacity at residual boutons. Sall3^Het^ mice displayed intermediate reductions across these same measures, likely reflecting partial loss of Sall3 expression in DDH inhibitory interneurons (Figure 2I).

The reduction in GABApre boutons was not secondary to loss of proprioceptive sensory afferents or a general disruption of spinal inhibitory synapses. vGlut1^+^ terminal number was only modestly increased in the VH of Sall3^cKO^ mice (Figures 3E,J; Figure S4D), likely reflecting increased sampling density associated with the smaller overall size of Sall3^cKO^ mice (Figure S3A). To assess inhibitory synapses more broadly, we used GAD67 (G67) to label GABApre boutons in addition to GABApost synapses, which contact motor neurons and interneurons.^6,12^ GABApre contacts were similarly depleted when visualized with G67 (Figure S4I), whereas GABApost contacts were unchanged in Sall3^Gad2-cKO^ mice (Figure S4J). Together, these results identify GABApre boutons as the principal circuit element disrupted by Sall3 loss.

To determine if the loss of G65^on^ boutons reflects decreased presynaptic G65 expression or loss of the GABApre terminal itself, we crossed *Sall3* conditional mice to a *Gad2*^GFP^ reporter, enabling direct visualization of *Gad2*^+^ axons and terminals. Sall3^GFP-Ctl^ mice exhibited dense GFP^+^ projections throughout the ventral horn, including contacts onto vGlut1^+^ terminals. In contrast, Sall3^GFP-cKO^ mice showed a striking depletion of GFP^+^ axons in the ventral horn (Figure 3I). Using a masking strategy to isolate GFP^+^ contacts on vGlut1 terminals (GFP^on^; Figure S4H), GFP^on^ puncta were reduced by 89.4 ± 3.5% (v1) and 70.5 ± 6.8% (v2) (Figure 3J), and the mean surface area of remaining GFP^on^ puncta was reduced by 30-45% (Figure 3K). Although the reduction in GFP^on^ contacts was less pronounced than that observed for G65^on^ puncta, particularly in v2, this discrepancy likely reflects inclusion of passing GFP^+^ axons in the masking approach, consistent with the greater total area of GFP^+^ signal in v2 relative to v1 (Figure S4G).

To distinguish developmental and maintenance functions of Sall3, we next asked whether established GABApre boutons require continued Sall3 expression. To delete Sall3 in mature mice, we used an inducible *Gad2^CreER^* approach in which tamoxifen was administered at P25-P60, after GABApre circuits are established (Figure 4A). This resulted in a significant but partial loss of G65^on^ puncta, with puncta number reduced by 36.5 ± 11.2% (v1) and 51.5 ± 10.5% (v2). The surface area of remaining G65^on^ boutons was also reduced by 40-45%, and fluorescence intensity was decreased (Figures 4B-E). Although residual boutons likely reflect incomplete recombination, the magnitude of G65^on^ bouton loss indicates that Sall3 is required to sustain GABApre boutons in the adult spinal cord.

**Figure 4.**
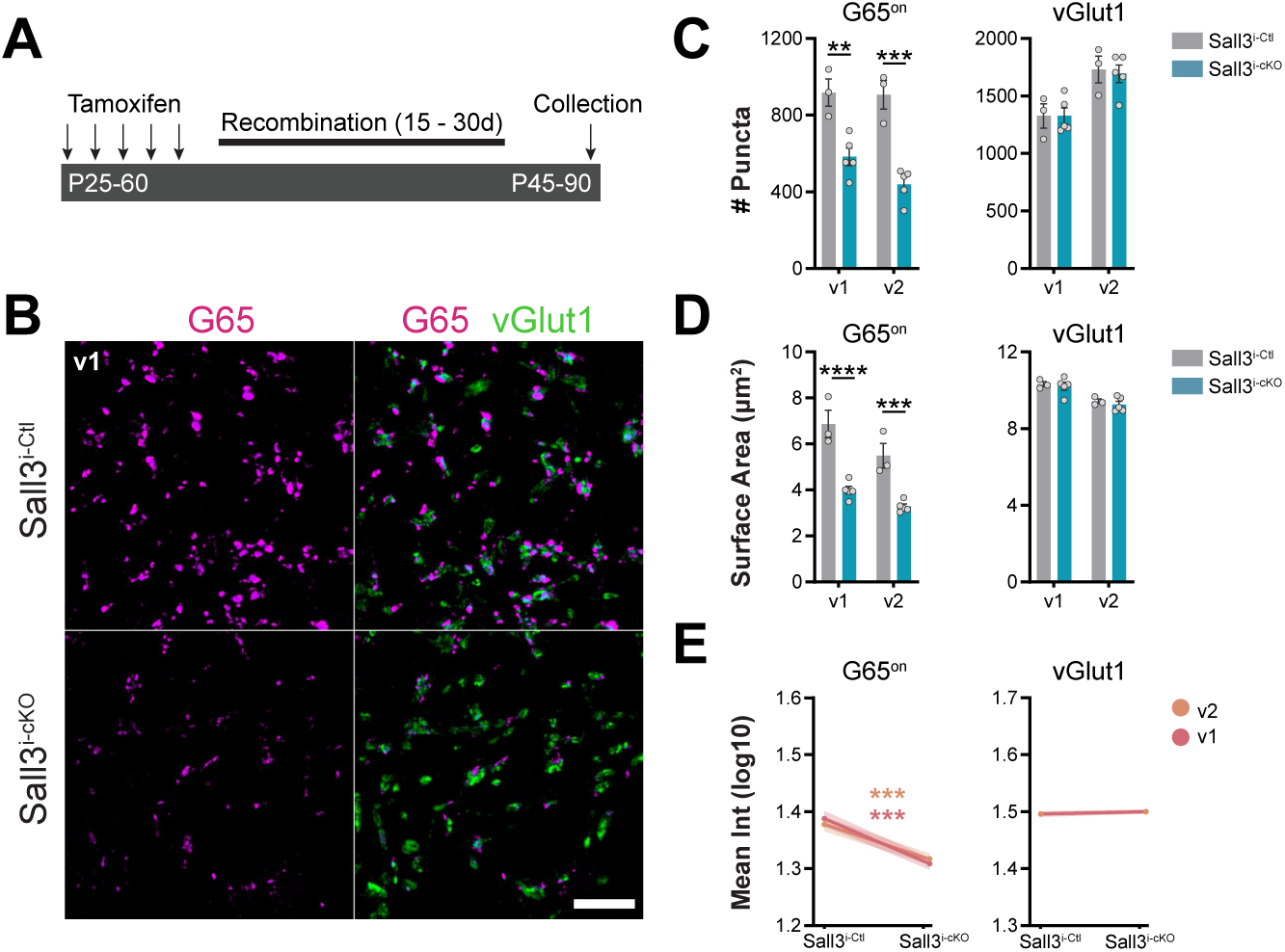
Sall3 is required for the maintenance of GABApre boutons in postnatal mice. **(A)** Schematic of inducible deletion of *Sall3* in mature GABAergic interneurons using the Gad2^CreER^ driver line. Tamoxifen was administered between P25 and P60 for five consecutive days, followed by a 15-30 day recombination period prior to analysis. **(B)** Immunostaining for G65 (GAD65, magenta) and vGlut1 (green) in the v1 region of the ventral spinal cord in representative Sall3^i-Ctl^ (top) and Sall3^i-cKO^ (bottom) mice. **(C-D)** Quantification of synapse reconstructions following tamoxifen-mediated *Sall3* deletion. The number of puncta per section (C) and mean surface area (D) are shown for G65^on^ and vGlut1. **(E)** Mean fluorescence intensity (Int) of reconstructed G65^on^ and vGlut1 surfaces plotted as a function of gene dose with fitted trends. Scale bars: (B) 10 µm. Graphs show mean ± SEM, with dots representing individual mice. Statistical significance is indicated as ** *p* < 0.01, *** *p* < 0.001, **** *p* < 0.0001. Exact *p* values, sample sizes (N), means, and standard deviations are provided in Table S1.

### Loss of GABApre boutons in Sall3^cKO^ mice disrupts presynaptic inhibition of sensory transmission

To determine how loss of GABApre boutons alters sensory transmission within spinal reflex circuits, we examined monosynaptic reflex (MSR) function in an *ex vivo* adult sacral spinal cord preparation across three distinct stimulation paradigms. We first established baseline MSR responses by stimulating the S4 dorsal root (DR) and recording the ventral root reflex (VRR) from the corresponding S4 ventral root (VR), quantified as the peak-to-peak amplitude of the compound action potential (CAP). In parallel, we monitored primary afferent depolarization (PAD) from a neighboring DR (S2), measured as the peak amplitude of the evoked dorsal root potential. This PAD signal serves as a readout of afferent excitability rather than a direct measure of presynaptic inhibition (Figure 5A).^8,9^ PAD and VRR increased with progressive activation of sensory afferents, consistent with expected recruitment dynamics (Figure 5B; Figures 5C,D, left).^2,43^ Subsequent analyses focused on low (1.5×T) and high (10×T) stimulation intensities, corresponding predominantly to Ia-mediated and broader cutaneous/polysynaptic recruitment regimes, respectively. At 1.5×T, PAD amplitude was comparable between control and Sall3^cKO^ mice, whereas VRR amplitude was reduced in Sall3^cKO^ mice (Figures 5C,D, middle). In contrast, at higher stimulation intensity (10×T), both PAD and VRR responses were diminished in Sall3^cKO^ mice relative to controls (Figures 5C,D, right).

**Figure 5.**
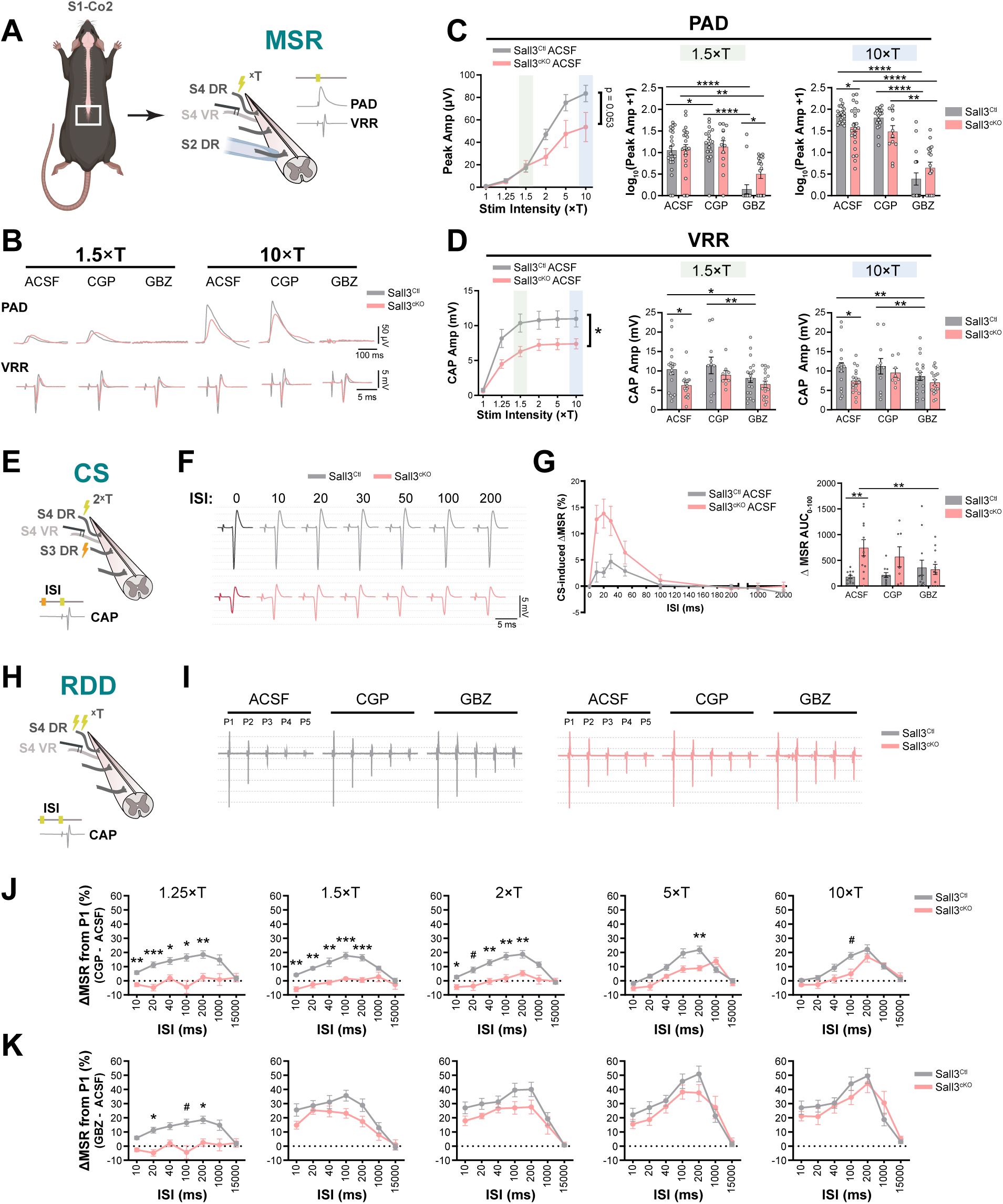
GABA_A_-mediated PAD and GABA_B_-mediated presynaptic inhibition are disrupted in Sall3^cKO^ mice. **(A)** The sacrocaudal spinal cord (S1–Co2) was isolated for *ex vivo* electrophysiological recordings of primary afferent depolarization (PAD) and ventral root reflexes (VRRs). The S4 dorsal root (DR) was stimulated at varying intensities (×T), while PAD was recorded from the S2 DR using a suction electrode and motor output (compound action potential, CAP) was recorded from the S4 ventral root (VR). **(B)** Representative PAD (top) and VRR (bottom) traces from Sall3^Ctl^ and Sall3^cKO^ mice recorded during baseline ACSF, GABA receptor blockade (CGP55485; CGP), or GABA_A_ receptor blockade (gabazine; GBZ) at 1.5×T (left) and 10×T (right) stimulation intensities. **(C)** Quantification of PAD responses. Left: peak amplitude across stimulus intensities under baseline ACSF conditions. Middle and Right: PAD responses at 1.5×T and 10×T stimulation, respectively, under ACSF, CGP, and GBZ conditions. **(D)** Quantification of VRR responses. Left: CAP amplitude across stimulus intensities under baseline ACSF conditions. Middle and Right: CAP responses at 1.5×T and 10×T stimulation, respectively, under ACSF, CGP, and GBZ conditions. **(E)** Experimental setup for conditioned-stimulus (CS) recordings. A conditioning stimulus was delivered to the S3 DR, followed by a test stimulus to the S4 DR at varying interstimulus intervals (ISI). Motor output was recorded from the S4 VR. **(F)** Representative test-evoked CAP responses from Sall3^Ctl^ (top) and Sall3^cKO^ (bottom) mice at ISIs of 0, 10, 20, 30, 50, 100, and 200 ms under ACSF baseline conditions. **(G)** Quantification of test-evoked monosynaptic reflex (MSR) responses. Left: CS-induced ΔMSR at each ISI, expressed as the percent change from the corresponding unconditioned response under baseline ACSF conditions. Right: area under the ISI-response curve from 0-100 ms (AUC_0-100_) under ACSF, CGP, and GBZ conditions. **(H)** Experimental setup for rate-dependent depression (RDD) recordings. The S4 DR was stimulated repeatedly with five pulses at varying ISIs, and output was recorded from the S4 VR. **(I)** Representative RDD traces from Sall3^Ctl^ and Sall3^cKO^ mice showing depression over five consecutive pulses (P1-P5). Traces are displayed with comparable P1 amplitudes across genotypes and conditions to facilitate comparison of subsequent pulse responses independent of initial response differences. **(J-K)** Effects of CGP (J) and GBZ (K) on RDD are plotted across ISIs and stimulus intensities (1.25-10×T). MSR responses were normalized to the first pulse (P1), averaged across pulses P2-P5, and expressed as percent change from P1. Drug effects are shown as the change relative to ACSF (drug − ACSF). Graphs show mean ± SEM, with dots representing individual hemicord recordings analyzed as independent samples. Additional representative traces and quantifications for drug conditions are provided in Figures S5A,C. Statistical significance is indicated as * *p* < 0.05, ** *p* < 0.01, *** *p* < 0.001, **** *p* < 0.0001. # indicates 0.05 ≤ *p* < 0.07. Exact *p* values, sample sizes (N), means, and standard deviations are provided in Table S1.

To examine the relative contributions of GABA_A_ and GABA_B_ to PAD and reflex transmission, we applied receptor-specific antagonists. Across stimulation intensities, gabazine (GBZ) abolished PAD responses in both genotypes, consistent with the established GABA_A_-dependence of PAD.^7,9,12,24^ In contrast, GABA_B_ receptor blockade (CGP) modestly increased PAD in controls at low stimulation intensities but had little effect in Sall3^cKO^ mice or at higher stimulation intensities in either genotype (Figure 5C; Figures S5A,B). For VRR responses, the effects of GBZ and CGP were similar across stimulation intensities. GBZ modestly reduced VRR amplitude in controls but had little to no effect in Sall3^cKO^ mice, whereas CGP had no effect in either genotype (Figure 5D; Figures S5C,D). At near-threshold stimulation (1×T), pharmacological manipulation unmasked weak MSR responses that were not apparent under baseline conditions (Figure S5E). Both CGP and GBZ increased reflex amplitude in controls, while only GBZ increased the reflex in Sall3^cKO^ and CGP had little effect. This suggests that inhibitory mechanisms strongly constrain sensory transmission near threshold, with a GABA_B_-dependent component that is diminished in Sall3^cKO^ mice.

To examine the relationship between presynaptic depolarization and motor output, we compared PAD and VRR amplitudes at low or high stimulation intensities (Figure S5F). PAD and VRR were not significantly correlated in either genotype at low stimulation intensities. However, at high stimulation, PAD and VRR were positively correlated in controls but negatively correlated in Sall3^cKO^ mice, indicating a disruption in the relationship between afferent depolarization and reflex output.

As a second stimulation paradigm, we employed a conditioning stimulus (CS) approach, which is widely used to assess presynaptic inhibition.^9,12,22,44–46^ In this paradigm, activity in one sensory pathway suppresses transmission in another, enabling measurement of heterosynaptic modulation within spinal reflex circuits. Accordingly, we delivered a CS to the S3 DR followed by a test stimulus to the S4 DR (Figure 5E). Because a strong CS can produce PAD-driven postactivation depression that may be mistaken for presynaptic inhibition,^45^ we set the CS intensity below the threshold required to evoke an S4 ventral root response. At each interstimulus interval (ISI), CS-induced ΔMSR was calculated as the percent change in the test-evoked MSR relative to the corresponding unconditioned response. Under baseline ACSF conditions, both genotypes showed brief facilitation at short ISIs (Figures 5F,G; Figure S5G), indicating transiently enhanced sensory transmission. This facilitation was markedly larger in Sall3^cKO^ mice, with the area under the curve over the 0-100 ms ISI window (AUC_0-100_) approximately three times that of control mice (Figure 5G, right). Blocking GABA_B_ receptors with CGP had no effect in either genotype, whereas GABA_A_ receptor blockade with GBZ abolished the enhanced facilitation in Sall3^cKO^ mice, indicating this phenotype depends on GABA_A_-mediated signaling.

As a final stimulation paradigm, we examined rate-dependent depression (RDD), a homosynaptic form of plasticity in which repeated activation of a single sensory pathway progressively reduces reflex amplitude.^1,47^ Five-pulse stimulus trains were delivered to the S4 DR, and MSR responses in the S4 VR were measured and expressed as the percent change from the first pulse (P1; Figures 5H,I). The overall magnitude of depression was comparable between genotypes under baseline ACSF conditions, but pharmacological dissection revealed distinct underlying mechanisms contributing to RDD (Figure S5H). Blockade of GABA_A_ receptors with GBZ partially relieved RDD in both genotypes across most stimulus intensities. At low stimulus intensity (1.25×T), however, GBZ produced a greater relief of depression in control mice (∼34% increase) compared to Sall3^cKO^ mice (∼19% increase; Figure 5J). In contrast, GABA_B_ receptor blockade with CGP revealed a pronounced genotype-specific difference. At low to moderate stimulus intensities (1.25-2×T), CGP relieved RDD in the control mice (∼15-20% increase) but had little effect in Sall3^cKO^ mice (∼1-2% increase; Figure 5K). At higher stimulus intensities, however, CGP produced a similar relief of RDD in both genotypes, eliminating the genotype difference observed at lower intensities and indicating that the GABA_B_-dependent component of RDD during predominantly Ia-driven recruitment is lost in Sall3^cKO^ mice.

### Loss of GABApre boutons impairs fine motor coordination and proprioceptive-dependent behaviors

To determine how the disruption of presynaptic inhibition impacts motor behavior, we evaluated Sall3^cKO^ mice using locomotor and skilled motor tasks. We first assessed baseline locomotion and gait using the CatWalk XT system (Figure 6A). Stride length, step cycle duration, and duty cycle were comparable across genotypes (Figures S6F-H), indicating gross locomotor rhythm was preserved.^48,49^ However, subtle differences in stance and limb coordination were present. Sall3^cKO^ mice exhibited reduced hindpaw contact area with the runway and a widened hindlimb base of support (Figures 6B,C). Analysis of step sequence patterns revealed reduced cruciate alternating/bound (CA/B) and alternate (AA) patterns, accompanied by an increase in abnormal alternating patterns (AB; Figure 6D). In addition, diagonal interlimb phase relationships were reduced (Figures S6A-E), consistent with weakened diagonal coupling. To complement gait analysis, we performed unsupervised behavioral profiling using Motion Sequencing (MoSeq) revealed a shift in the locomotor syllable repertoire, with Sall3^cKO^ mice favoring lower-velocity movement motifs during active locomotion (Figures S6I-O). Together, these findings indicate that Sall3^cKO^ mice have mild coordination deficits underlying otherwise normal gait.

**Figure 6.**
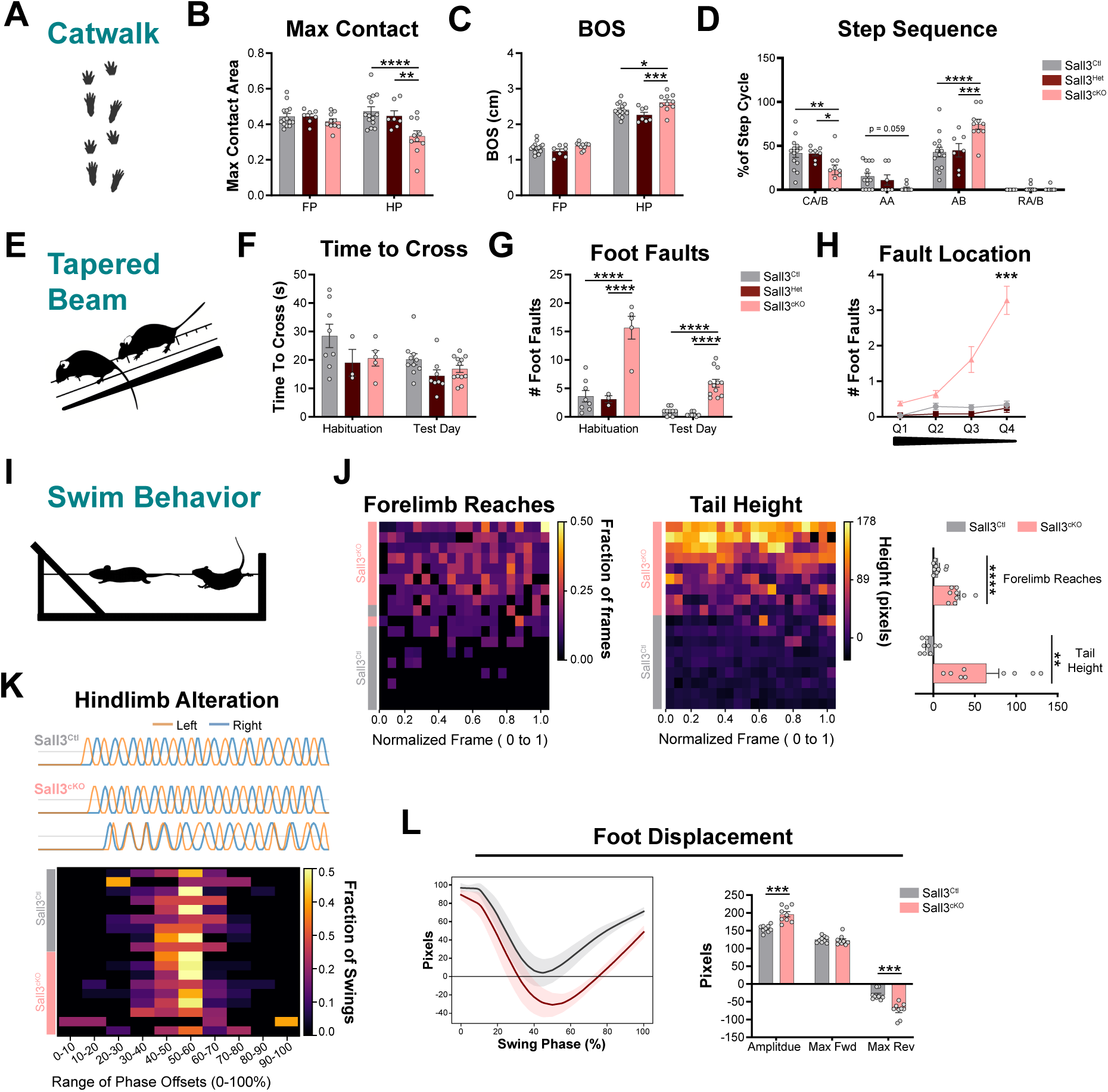
Altered presynaptic inhibition impairs fine motor control and swim behavior. **(A)** Illustration of the CatWalk XT gait analysis system. Mice traverse an illuminated glass runway while paw contacts are recorded by a high-speed camera positioned beneath the walkway. **(B-C)** Quantification of CatWalk parameters in Sall3^Ctl^, Sall3^Het^, and Sall3^cKO^ mice. Maximum contact area for FP and HP on the glass runway (B) and distance between left and right paws of the same limb pair (BOS, C). **(D)** Step sequence patterns describing the order of paw placement during locomotion. **(E)** Illustration of the tapered beam assay. The beam narrows progressively as mice traverse toward an enclosed goal box. **(F-H)** Quantification of tapered beam performance during habituation (first exposure) and the subsequent test day. Time to cross (F), number of foot faults (G), and spatial distribution of foot faults along the beam during the test day (H). **(I)** Illustration of the swim assay. Mice swim across a narrow water-filled tank toward an escape platform. **(J)** Swimming forelimb and tail movements across normalized swim duration (0-1), binned into 20 equal intervals. Left: Heatmap of forelimb activity. Values represent the fraction of frames within each bin in which either forelimb was engaged. Middle: Heatmap of vertical tail-tip height relative to the nose, with each bin showing the maximum value. Right: Per-mouse quantification of total forelimb reaches and mean maximum tail height across bins. **(K)** Quantification of hindlimb alternation. Top: Representative swing phase traces from Sall3^Ctl^ (gray), Sall3^cKO^ mice with alternating patterns (top), and Sall3^cKO^ mice with synchronous patterns (bottom). Left and right hindlimbs are shown in orange and blue, respectively. Bottom: Heatmap showing the fraction of hindlimb swings within each phase-offset bin across mice. Values of 0.5 indicate alternating limb movement, whereas values of 0 or 1 indicate synchronous movement. **(L)** Quantification of hindlimb foot displacement during swing. Left: Phase-normalized displacement traces (0-1) averaged across swings. Right: Quantification of average swing amplitude, Max Fwd, and Max Rev. Graphs show mean ± SEM, with dots representing individual mice. Statistical significance is indicated as \**p* < 0.05, \*\**p* < 0.01, \*\*\**p* < 0.001, \*\*\*\**p* < 0.0001. In (H), *** denotes both the Sall3^Ctl^ vs Sall3^cKO^ and Sall3^Het^ vs Sall3^cKO^ comparisons. Exact *p* values, sample sizes (N), means, and standard deviations are provided in Table S1. Abbreviations: FP, front paws; HP, hind paws; BOS, base of support; CA/B, cruciate alternating/bound; AA, alternate; AB, abnormal alternating; RA/RB, rotary; Max Fwd, maximum forward displacement; Max Rev, maximum reverse displacement.

To evaluate whether additional motor deficits emerge under increased task demands, we tested Sall3^cKO^ mice using assays requiring coordination and balance. Control and Sall3^cKO^ mice performed comparably on an accelerating rotarod (Figure S6P), indicating that gross motor coordination and endurance are preserved in Sall3^cKO^ mice. We next assessed precision stepping and adaptive balance using a tapered beam that progressively narrows along its length (Figure 6E). Despite similar overall traversal times between genotypes (Figure 6F), Sall3^cKO^ mice exhibited a marked increase in foot faults on both habituation and test days (habituation: 15.7 ± 4.5 vs 3.6 ± 2.8; test: 5.9 ± 2.4 vs 0.9 ± 0.7; Figure 6G, Video S1). This deficit persisted despite improved performance across days, particularly on the narrowest beam segments (Figure 6H).

Loss of presynaptic inhibition has been linked to altered goal-directed forelimb behavior in pellet-reaching tasks,^12^ so we next assessed forelimb function using a similar assay (Figure S6Q). Sall3^cKO^ mice achieved 2.5-fold fewer successful pellet retrievals than controls (Sall3^Ctl^: 18.8 ± 10.4; Sall3^cKO^: 7.6 ± 6.9). However, control mice performed below expected levels and failed to show robust improvement over the 10-day training period, limiting the interpretation of the magnitude of this deficit. When Sall3^cKO^ mice initiated reaches, movements appeared smooth and lacked obvious abnormalities (Video S2).

To specifically examine motor control under conditions where load-dependent compensatory input is reduced,^50^ we next evaluated swimming behavior in Sall3^cKO^ mice (Figure 6I). Control mice maintained largely stationary forearms with minimal tail movement. In contrast, Sall3^cKO^ mice recruited their forelimbs more frequently (Sall3^Ctl^: 5.2 ± 6.1; Sall3^cKO^: 28.7 ± 10.1) and exhibited increased tail elevation (Sall3^Ctl^: -6.7 ± 7.8 pixels; Sall3^cKO^: 63.8 ± 44.8 pixels) during propulsion (Figure 6J, Video S3). Hindlimb phase relationships remained largely intact, with most Sall3^cKO^ mice exhibiting alternating hindlimb activation, although occasional mice displayed phase-offset distributions consistent with synchronous activation (Figure 6K). In contrast, analysis of hindlimb swing kinematics revealed a robust motor phenotype (Figure 6L). Early swing trajectories were similar between genotypes, but overall swing amplitude (Sall3^Ctl^: 163.2 ± 16.9; Sall3^cKO^: 195.1 ± 23.1) and maximal reverse displacement (Sall3^Ctl^: -38.2 ± 16.7; Sall3^cKO^: -73.7 ± 22.2) were increased in Sall3^cKO^ mice, consistent with hindlimb overextension during propulsion. Cumulatively, these motor phenotypes indicate that GABApre boutons are essential for refining sensory-motor integration under increased task demands that require precise coordination and regulated proprioceptive feedback.

## Discussion

The present study identifies Sall3 as a principal determinant of GABApre inhibitory circuitry. Loss of Sall3 results in elimination of GABApre boutons from proprioceptive Ia afferent terminals, impaired presynaptic regulation of sensory transmission, and deficits in motor behaviors that rely on accurate proprioceptive feedback. Moreover, inducible deletion of Sall3 in adulthood produced substantial GABApre terminal loss, indicating that Sall3 is required not only during circuit assembly but also for the long-term maintenance of mature presynaptic inhibitory synapses.

The selective loss of GABApre terminals in Sall3 conditional knockout mice provided an opportunity to dissect the contributions of terminal presynaptic inhibition to sensory transmission throughout the spinal cord. Despite the absence of GABApre boutons, PAD evoked by low-threshold dorsal root stimulation remained largely intact in Sall3^cKO^ mice. Moreover, conditioning stimulation of a neighboring dorsal root produced greater facilitation of sensory transmission than in littermate controls, an effect abolished by the GABA_A_ receptor antagonist gabazine. Thus, Sall3-dependent GABApre boutons act to limit the facilitatory effect of GABA_A_-dependent PAD on afferent transmission.^9,51^

This interpretation creates an important physiological tension. If the major consequence of losing GABApre terminals was simply reduced inhibition of transmitter release, monosynaptic ventral root reflexes would be expected to increase. Instead, reflex amplitudes were consistently reduced in Sall3^cKO^ mice. The relationship between PAD and reflex output was also markedly altered, indicating that Sall3 deletion disrupts the normal coupling between afferent depolarization and reflex output. Although the basis for this paradoxical phenotype remains unclear, it may reflect compensatory reorganization of spinal circuits in response to developmental loss of presynaptic inhibition.

A clearer role for Sall3-depenent presynaptic inhibition emerged from experiments probing GABA_B_ receptor function. Blockade of GABA_B_ receptors in controls increased ventral root reflex amplitudes evoked by stimulation at threshold and relieved homosynaptic rate-dependent depression but had little effect in the absence of Sall3, indicating that tonic and activity-dependent GABA_B_-mediated regulation of sensory transmission are lost following Sall3 deletion. Together, these findings support an emerging model in which GABA_A_-mediated signaling promotes PAD and facilitates spike propagation at sensory axon nodes, whereas Sall3-dependent GABApre boutons mediate GABA_B_-dependent presynaptic inhibition that limits transmitter release from proprioceptive Ia terminals.^9,25,45^

The behavioral effects of Sall3 deletion closely mirror those anticipated following disruption of Ia presynaptic inhibition. Sall3^cKO^ mice maintained normal locomotor rhythm and gross gait parameters during overground locomotion, although subtle changes in postural and interlimb control hinted that compensatory mechanisms might be masking an underlying motor deficit. As behavioral tasks placed greater demands on proprioceptive feedback, the effects of Sall3 deletion on motor behavior became apparent. Swimming is particularly sensitive measure of proprioceptive function because the absence of load-related and contact-dependent sensory cues increases reliance on muscle spindle-derived proprioceptive feedback.^50^ Under these conditions, Sall3^cKO^ mice lost normal phase-dependent modulation of hindlimb movement and resorted to increased forelimb recruitment and exaggerated tail movements. Skilled beam walking uncovered another dimension of this phenotype. Sall3^cKO^ mice could walk across the wider portions of the tapered beam but made more foot placement errors with decreasing beam width. However, in keeping with previous evidence that reflex pathways can be modified through training and experience,^10,52^ foot placement accuracy increased with repeated testing. Collectively, these behavioral findings suggest that Sall3-dependent presynaptic inhibition mediates rapid, context-dependent gain control of Ia sensory transmission during movement.

Our behavioral findings extend previous work by Fink and colleagues,^12^ who used genetic ablation of *Gad2*-expressing presynaptic inhibitory neurons to investigate the role of GABApre circuitry in motor control. Like our study, Fink et al. found that disrupting presynaptic inhibitory circuitry produced selective impairments in sensorimotor behavior. However, several experimental differences are important to consider when comparing their findings with ours. Fink et al. did not report locomotor deficits, instead their behavioral analysis emphasized a role for Gad2-expressing presynaptic inhibitory neurons in goal-directed reaching. Although Sall3^cKO^ mice were also less successful during goal-directed reaching, they did not exhibit the prominent limb oscillations or exaggerated corrective movements reported by Fink et al. This could reflect differences in the genetic manipulations themselves, with *Gad2*-driven ablation affecting a broader neuronal population, as suggested by the accompanying itch phenotype, or the acute disruption of a mature circuit in Fink et al. compared with the lifelong loss of presynaptic inhibition in Sall3^cKO^ mice. Importantly, our study enabled a more direct assessment of hindlimb locomotor function. Whether Fink et al. did not detect a locomotor phenotype because the horizontal ladder places different demands on foot placement than the tapered beam, or because their manipulation was restricted to the cervical spinal cord segments where descending pathways could provide additional compensation, remains unclear. Because the tapered beam phenotype in Sall3^cKO^ mice was driven almost entirely by hindlimb foot placement errors, the latter explanation may be particularly relevant. This underscores the value of a spinal cord-wide disruption of GABApre terminals for revealing locomotor deficits that may not be apparent with more restricted manipulations.

Sensory afferents terminate within anatomically distinct domains of the spinal cord, with proprioceptive and cutaneous inputs occupying different dorsoventral region,^53^ and accumulating evidence suggests that multiple presynaptic inhibitory interneuron subtypes target specific sensory feedback pathways.^12^ The dorsoventral gradient of presynaptic inhibitory bouton loss following Sall3 deletion supports this model, suggesting that Sall3 defines only a subset of presynaptic inhibitory interneurons. Within proprioceptive circuits muscle spindle (Ia and II) and Golgi tendon organ (Ib) afferents terminate in the ventral and deep dorsal horn or predominately in the deep dorsal horn, respectively.^54^ The complete loss of GABApre boutons in the ventral horn but only a partial reduction in the deep dorsal horn aligns with the idea that Sall3-associated presynaptic inhibitory interneurons preferentially target muscle spindle afferents, while the remaining deep dorsal horn boutons potentially arise from a distinct Ib-associated presynaptic inhibitory population.

One candidate for this remaining population are the Rorβ-derived presynaptic inhibitory interneurons,^20^ which does not project to the ventral horn (Figures S2J,K) and produces a ‘hopping gait’ locomotor phenotype distinct from that observed following Sall3 deletion ^20^. Similarly, the partial preservation of presynaptic inhibitory boutons in the dorsal horn suggests that Sall3 contributes to only a subset of cutaneous presynaptic inhibitory interneurons. Several genetically defined presynaptic inhibitory interneuron populations have been identified in this region, including parvalbumin-expressing interneurons that regulate myelinated cutaneous afferents and calretinin-expressing interneurons associated with non-peptidergic nociceptors.^18,19^ Whether Sall3 contributes to one of these established populations or instead identifies a previously unrecognized presynaptic inhibitory interneuron population remains to be determined.

The abundance of Sall3-expressing inhibitory interneurons within the dorsal horn predicts that presynaptic inhibition is only one component of Sall3 function. Sall3^cKO^ mice developed pathological itch and exhibited enhanced chemically evoked itch despite largely preserved tactile, thermal, and acute nociceptive responses. Although Sall3 is largely excluded from the mature dynorphin interneurons that suppress chemical itch,^55,56^ it is transiently co-expressed with Bhlhe22 during development in the population that gives rise to these neurons (Figures S3K,L).^57^ This transient developmental relationship, together with the loss of dorsal Pax2^+^ inhibitory neurons in Sall3^cKO^ and Bhlhe22 mutant mice, raises the possibility that Sall3 contributes to the Bhlhe22-dependent developmental program. However, unlike Bhlhe22 mutants,^57^ pathological itch in Sall3^cKO^ mice was less severe and incompletely penetrant, and persistent inflammatory pain responses during phase II of the formalin test were reduced rather than enhanced. Because both behavioral phenotypes have been proposed to arise as downstream consequences of chronic disinhibition of chemical itch pathways, the contrasting phenotype observed in Sall3^cKO^ mice suggests a more complex relationship between inhibitory itch circuits and complementary pathways involved in pain processing and sensitization.

Together, these findings establish Sall3 as a molecular entry point for understanding inhibitory circuits that regulate sensory processing across the spinal cord, from Ia presynaptic inhibition to broader dorsal inhibitory networks involved in pain and itch. Future intersectional genetic approaches combining Sall3 with complementary molecular markers should enable these populations to be dissected individually, revealing how specialized inhibitory circuits shape sensory transmission and behavior.

### Limitations of the Study

Although we identify a central role for Sall3 in spinal inhibitory circuitry, two important questions remain unresolved. While inducible deletion of Sall3 in adulthood eliminated about half of GABApre boutons, this loss was less extensive than following developmental deletion. Because *Gad2^CreER^*-mediated recombination was incomplete, it remains unclear whether partial GABApre bouton loss reflects a reduced requirement for Sall3 in the mature circuit or incomplete targeting of Sall3-expressing interneurons. In addition, although the behavioral and electrophysiological findings are most consistent with a primary spinal mechanism, Sall3 is also expressed in a small number of discrete GABAergic populations in the brain, as identified using the Allen Brain Atlas (Table S2).^58,59^ Although this restricted expression argues against widespread supraspinal involvement, contributions from these populations cannot be excluded. Future studies combining more efficient adult deletion and spinal-cord restricted genetic strategies will therefore be important for defining the specific temporal and anatomical roles of Sall3 in inhibitory circuit function.

## Materials and Methods

### Animals

All animal procedures were conducted in accordance with the NIH Guidelines for the Care and Use of Laboratory Animals. Experiments performed at Stanford University were approved by Stanford’s Administrative Panel on Laboratory Animal Care (APLAC), and electrophysiology experiments performed at Northwestern University were approved by the Northwestern University Animal Care and Use Committee. Mice of both sexes on a C57BL/6 background were used throughout this study. Animals were group-housed, up to five adults per individually ventilated cage, in a temperature- and humidity-controlled facility with food and water provided *ad libitum*. Most experiments were performed on mice maintained on a standard 12-hour light/dark cycle; however, animals used for MoSeq, rotarod, and CatWalk assays were housed on a reverse light cycle. Embryonic analyses were carried out at E12.5 and E13.5, neonatal studies at P0 and P7, and synaptic analyses at P25. Conditional deletion of *Sall3* in adulthood was assessed after 1.5-3 months of age, and behavioral experiments were performed in adult mice between 3-7 months old.

#### Mouse Lines

All mice used for this study were either purchased or acquired as detailed in the table below. The following genotypes were generated and used: *Ptf1a^Cre/Cre^*(Ptf1a^KO^) and *Ptf1a^+/+^* (Ptf1a^Ctl^);^60^ *Gad2^Cre/+^*;*Sall3^fx/fx^*(Sall3^cKO^), *Gad2^Cre/+^*;*Sall3^fx/+^* (Sall3^Het^), and *Gad2^+/+^*;*Sall3^fx/fx^* or *Gad2^+/+^*;*Sall3^fx/+^*(pooled as Sall3^Ctl^). To visualize *Gad2*-expressing projections in *Sall3* conditional mutants, we incorporated the *Gad65^N^*^45^*^-GFP^* transgenic reporter, in which GFP expression is driven by *Gad2* (GAD65) regulatory sequences,^61^ generating *Gad2^Cre/+^*;*Sall3^fx/fx^*;*Gad65^N^*^45^*^-GFP/+^* (Sall3^GFP-cKO^) and corresponding control littermates (Sall3^GFP-Ctl^) analogous to those described for *Gad2^Cre^*. We also used *Pvalb*^tdTomato/+^ (PV^tdT^) and *Rorb^Cre/+^*;Ai14^tdT/+^ (Rorb^tdT^) mice for lineage tracing experiments and generated *Gad2^CreER/+^*;*Sall3^fx/fx^* (Sall3^i-cKO^) and *Gad2^+/+^*;*Sall3^fx/fx^* (Sall3^i-Ctl^) to assess the role of Sall3 in adult mice.

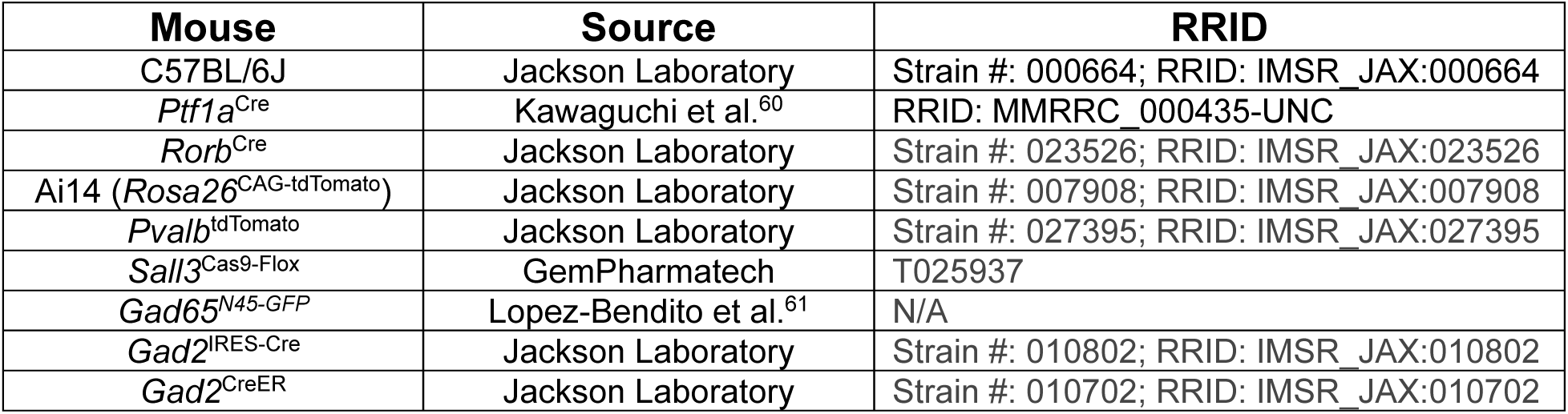

#### Sall3-Conditional Line

The Sall3 conditional allele (*Sall3^Cas9-Flox^*) was generated by GemPharmatech using CRISPR/Cas9-mediated insertion of loxP sites flanking exon 2 and exon 3 of the *Sall3* gene. CRISPR/Cas9 reagents and the donor construct were microinjected into fertilized C57BL/6JGpt zygotes and transplanted to obtain F0 founders. F0 positive mice were bred to C57BL/6JGpt animals to produce an F1 generation, and F1 carriers were intercrossed to establish a stable homozygous line. *Sall3* genotyping was performed using 5’-arm (T025937-F1: TGCAAGCACCAAGAAGTATCACCG and T025937-R1: CATGAGAATGCTGTAGTCAGACACAC, yielding a 278-bp wild-type band or a 383-bp targeted band) and 3’-arm (T025937-F2: TGAGGTCTAGAATGTTCCACCGTCT and T025937-R2: GCTAGAGTCTCTTCCTGGCTTAGAACTTG, yielding a 439-bp wild-type or a 545-bp targeted band) primers.

#### Tamoxifen Treatment

Tamoxifen (Sigma-Aldrich, cat# T5648) was dissolved in corn oil at 20 mg/ml and vortexed for 1-2 hours to ensure complete solubilization. Adult mice received tamoxifen by oral gavage at a dose of 0.25 mg/g of body weight once daily for five consecutive days. Two independent cohorts were analyzed. In cohort 1, tamoxifen administration began at P25, and tissues were collected 15 days after the final dose. In cohort 2, tamoxifen administration began at P60, and tissues were collected 30 days after the final dose to allow for a longer interval for potential synaptic remodeling. Extending the post-induction interval did not increase the degree of GABApre bouton loss; therefore, data from both cohorts were pooled for analysis.

#### Timed Matings

Adult male and female *Ptf1a*^Cre/+^ mice were housed together overnight for timed matings. Females were checked each morning for the presence of a vaginal plug, and noon on the day a plug was detected was designated embryonic day 0.5 (E0.5). Plug-positive females were separated from males and maintained individually until tissue collection at the appropriate time points.

### Single-cell RNA-sequencing analysis

Publicly available embryonic cervical and thoracic spinal cord scRNA-seq data from^26^ were reanalyzed with a specific focus on identifying and characterizing the dI4/LA dorsal interneuron population. The Delile dataset includes two embryos each from E9.5, E10.5, and E11.5, and three embryos each from E12.5 and E13.5. Raw count matrices and metadata were merged into a single .h5ad object for analysis using Scanpy (v1.9.3).^62^ Doublets were detected on a per-embryo basis using Scrublet (v0.2.3)^63^ and predicted doublets were removed before downstream processing.

Low-quality genes and cells were filtered based on detection frequency, total counts (>1,000), number of expressed genes (>1,000), and abnormal mitochondrial (between 1 and 8%) or ribosomal RNA content (>8%). Counts were normalized to 10,000 reads per cell and log-transformed. Cell-cycle scores were computed using curated Regev lab gene sets.^64^ Highly variable genes (HVG) were selected (5,000), principal component analysis (PCA) was performed (n=20), and batch effects across embryos and developmental ages were corrected using BBKNN (v1.5.1).^65^ UMAP/umap-learn (v0.5.7)^66^ and Leiden clustering (v0.9.1)^67^ were computed from batch-corrected neighborhood data. Cell clusters were annotated using established marker sets for spinal progenitors, early neurons, DRG lineages, neural crest, mesoderm, interneurons, motor neurons, and non-neuronal lineages (see Figure S1F). To more precisely resolve the dI4/LA interneuron class, interneuron and dI4/LA clusters were subsetted and reclustered using a similar HVG-PCA-BBKNN-UMAP-Leiden workflow, and individual clusters were assigned to dI1-dI6 and V0-V3 lineages based on curated marker gene signatures (see Figure 1D). Differentially expressed genes were computed in Scanpy (rank_genes_groups, method=’t-test’). Genes were filtered using filter_rank_genes_groups requiring detection in ≥ 20% of dI4/LA cells, detection in ≤ 50% of V1 cells, and fold-change ≥ 1.5.

Genes enriched in the dI4/LA population from our embryonic analysis were compared to dI4/LA-enriched genes identified in Osseward et al.^33^ In that study, glutamatergic (*Vglut2^Cre^*;tdTomato) and GABAergic (*Vgat^Cre^*;tdTomato) spinal interneurons were FACS-purified from postnatal mice (three biological replicates per *Cre* line) and subjected to RNA-sequencing. Expression of candidate dI4/LA genes was queried directly from the authors’ published supplemental gene-expression tables, which provided their processed differential-expression results.

### Immunohistochemistry

#### Tissue Preparation

Mice were transcardially perfused with PBS to remove blood, and spinal cords were removed by hydraulic extrusion. Tissue was post-fixed in 4% paraformaldehyde (Electron Microscopy Sciences, cat# 15714) on ice for 2 hours, washed in PBS, and cryoprotected overnight in 30% sucrose. Lumbar spinal cords were dissected and divided into three segments corresponding approximately to L1-L2, L3-L4, and L5-L6. Segments were embedded in Tissue-Plus O.C.T compound (Fisher HealthCare, cat# 23-730-571) and frozen. L3-L4 segments were sectioned at 20 µm on a CM3050S cryostat (Leica Biosystems) across six serial slides such that sections on a given slide were about 120 µm apart.

#### Immunostaining

Cryosections were rehydrated in PBS (3 x 5 min) and incubated overnight at 4°C in primary antibodies (listed in the Key Resources Table) diluted in PBS containing 0.2% Triton X-100. The following day, sections were washed in 0.2% Triton X-100 (2 x 5 min) followed by PBS (1 x 5 min) and incubated for 2 hours at room temperature with donkey-derived secondary antibodies (Jackson ImmunoResearch) diluted in 0.2% Triton X-100. After final washes, slides were coverslipped with Fluoromount-G (SouthernBiotech).

| **Antibody** | **Source** | **Catalog #** | **RRID** |
| --- | --- | --- | --- |
| Rabbit anti-GAD65 (1:50,000) | Jessel Laboratory | CU1196 | AB_2904512 |
| Mouse anti-GAD67, clone 1G10.2 (1:10,000) | Sigma-Aldrich | MAB5406 | AB_2278725 |
| Chicken anti-GFP (1:1,000) | Millipore | AB16901 | AB_90890 |
| Mouse anti-NeuN, clone A60 (1:1,000) | Sigma-Aldrich | MAB377 | AB_2298772 |
| Goat anti-Pax2 (1:1,000) | R&D Systems | AF3364 | AB_10889828 |
| Rabbit anti-Sall3 (1:800) | Sigma-Aldrich | HPA016656 | AB_1856546 |
| Goat anti-RFP (1:1000) | OriGene | AB1140-100 | AB_2877097 |
| Guinea Pig anti-vGlut1 (1:5,000) | Jessel Laboratory | CU1327 |  |
| Donkey-derived secondary antibodies | Jackson ImmunoResearch | Assorted cat#s |  |

#### In Situ Hybridization

RNAscope Multiplex Fluorescent v2 assays (ACD Bio, cat# 323110) were performed according to the manufacturer’s instructions with minor modifications. Briefly, cryosections were baked at 60°C for 45 min, post-fixed in 4% paraformaldehyde for 60 min at room temperature, dehydrated through graded ethanol solutions (50%, 70%, 100%, 100%), and treated with hydrogen peroxide for 10 min. Antigen retrieval was performed at ∼95°C for 5 min using RNAscope Target Retrieval Reagent (ACD Bio), followed by an additional 45 min bake at 60°C. Sections were then incubated with Protease III for 40 min and hybridized with target probes (ACD Bio, *Pdyn*: cat# 318771-C3; *Sall3*: cat# 591791) for 2 h at 40°C in a HybEZ oven (ACD Bio, cat# 321710). Signal amplification and fluorescent detection were subsequently performed according to the manufacturer’s protocol. Slides were coverslipped with Fluoromount-G (SouthernBiotech).

#### Image Acquisition

Immunostained tissue was imaged on a Leica SP8 confocal microscope (Leica Biosystems). For cell-body analyses, images were acquired using a 20x objective (NA 0.75) at 2048 x 2048 resolution with a 2.5 µm z-step. Tile scans were collected using LAS Navigator and merged and max-projected in LAS X software (v1.4.7.28982). For synaptic analyses, images were acquired using a 63x objective (NA 1.40) at 3312 x 3312 resolution with a 0.3 µm z-step. Up to four non-overlapping regions were imaged per section: ventral horn region 1 (v1; most ventral), ventral horn region 2 (v2; slightly more dorsal/lateral), deep dorsal horn (DDH), and dorsal horn (DH) (see Figures 3A-C and Figures S4A-C).

#### Cell Body Quantification and Spatial Distribution Analysis

Maximum-intensity projections were imported into custom software, and cells within one hemisection of the spinal cord were manually annotated by an experimenter blinded to genotype and condition whenever possible. Regional boundaries were drawn within the software to assign each cell to its appropriate anatomical compartment. Cell identities (marker combinations) and XY coordinates were exported to Excel. Samples were decoded and cell counts were summarized per section and then averaged across sections to generate per-mouse values for each region. Counts were reported as raw values and as percentages of the total population within the same mouse and region. Three images were quantified per mouse.

To assess cell body spatial distribution, the center of the central canal in each section was used as a reference point to express all cell coordinates in a common anatomical frame. Coordinates were then oriented consistently across sections and normalized within each section to a unit square (0 – 1 range) to enable comparison across animals and developmental stages independent of tissue size. Spatial density was quantified using a two-dimensional kernel density estimation (KDE) on the normalized coordinates, computed separately for each mouse and cell group. Mouse-level density maps were averaged with equal weight per mouse to generate time point-level contours. KDE probability mass was integrated within predefined dorsoventral and mediolateral bins to yield per-mouse regional occupancy measures for statistical comparison across groups.

#### Synaptic Analysis

Imaris (Oxford Instruments, v9.8.2) was used to generate 3D surface reconstructions from confocal image stacks. Raw .lif files were imported and processed using batch mode to ensure identical segmentation parameters across images within each experiment, including background subtraction, median filtering (to smooth voxel intensity and reduce edge noise), intensity thresholding, and application of a minimum voxel cut-off. Two reconstruction pipelines were used depending on the analysis.

1. vGlut1-GAD65 analysis. vGlut1 surfaces were generated first. GAD65 surfaces were then reconstructed, and the Imaris shortest-distance metric was applied with a threshold of 0.05 µm to classify GAD65 signal in direct contact with a vGlut1 terminal as GAD65^on^.
2. vGlut1-GAD65-GFP and vGlut1-GAD67 analysis. In *Gad65*^N45-GFP^ mice and in analyses involving dense GAD67^+^ GABApost synapses, high signal density led to frequent surface merging and false-positive associations when reconstructing GFP or GAD67 objects directly. To restrict analysis to signal proximity to vGlut1 terminals, an adjusted workflow was implemented. vGlut1 surfaces were first reconstructed as described above. A distance-transformation dilation of 0.75 pixels was then applied to generate an expanded vGlut1 mask. This dilated vGlut1 object was used to mask the GFP or GAD67 channel prior to surface reconstruction. The masked GFP or GAD67 signal was subsequently reconstructed, and the shortest distance classifier was used to identify GFP^on^ (GABApre) or GAD67^off^ (GABApost) synapses (see Figure S4H for workflow).

Reconstruction output tables were exported from Imaris and per-mouse averages were calculated. For synaptic analyses, 8-10 images were quantified per region per mouse.

### Behavioral Assays

All behavioral assays were performed in adult mice between 3 and 7 months of age and were conducted by observers blinded to the genotype. Behavioral cohorts were run through multiple assays, with sufficient rest periods scheduled between tests to allow recovery and minimize carryover effects.

#### Body Weight

Mice were weighed at P1 and then every other day through P25. For each mouse, a linear regression was fit across time to calculate an individual growth rate (slope), which was used for statistical comparisons.

#### Dermatitis Evaluation

Spontaneous dermatitis was observed in a subset of Sall3^cKO^ mice. For animals enrolled in behavioral cohorts, the age at which dermatitis progressed to a severity requiring euthanasia was recorded, although these mice were not monitored continuously outside of behavioral testing schedules. Control littermates were euthanized when the final Sall3^cKO^ cage mate reached its dermatitis-defined endpoint.

#### Itch

Spontaneous itch was assessed by placing mice individually into clean, empty cages and observing them for 60 minutes. For chemical itch, histamine (10 µg/µL in saline; Sigma-Aldrich, cat# H7125) or chloroquine (10 µg/µL in saline; Sigma-Aldrich, cat# C6628) was injected intradermally (20 µL) at the nape of the neck, and mice were observed for 30 minutes, following established methods.^41^ In all assays, scratch bouts were scored throughout the observation period.

#### Pain

Capsaicin (2.5 µg in 20 µL; Sigma-Aldrich, cat# M2028) was prepared by dissolving capsaicin in 100% ethanol and diluting the solution so that the final injected concentration contained 2% ethanol. Formalin was prepared as 20 µL of a 2.5% solution made from 10% buffered formalin (Fisher HealthCare, cat# 23-305-510). Each solution was injected into the plantar surface of the hindpaw, following established procedures.^41^ Mice were observed for 10 minutes after capsaicin injection and for 60 minutes after formalin injection, and paw-licking behavior was scored during the observation periods.

#### Von Frey

Mice were placed on a 22-gauge wire mesh platform (McMaster-Carr, cat# 92725T2) inside clear plastic cylinders and acclimated for 60 minutes. Mechanical sensitivity was assessed using calibrated von Frey filaments (Bioseb, cat# BIO-VF-M) applied to the center of the hindpaw according to the up-down method.^68^ Testing began with the 3.22 g filament. If the mouse responded (paw withdrawal, licking, or jumping), an “X” was recorded; if no response occurred, an “O” was recorded. If the initial trial produced no response, progressively larger filaments were applied until a first positive response was obtained. After the first positive response, four additional stimulations were performed. After each stimulation, the filament size was adjusted according to the response, with positive responses followed by the next smaller filament and non-responses followed by the next larger filament. Seven to eight mice were tested in parallel during each trial block to allow adequate rest between trials. For each animal, the final filament applied (regardless of response) and the sequence of X-O responses were entered into a standard up-down method calculator to determine the withdrawal threshold.

#### Thermal

Cold and heat sensitivity were assessed following established procedures.^41^ Cold sensitivity was measured using a dry-ice evoked withdrawal assay. Mice were placed on a 4-mm tempered glass surface inside a plastic cylinder and acclimated for 10 minutes, after which dry ice was applied to the underside of the glass beneath the hindpaws until a withdrawal response was elicited (maximum of 2 minutes). Mice were rested for 15 minutes between trials, and data represent the average of three trials. Heat sensitivity was assessed using a hot plate, with mice placed inside a plastic cylinder on the heated surface for a maximum of 45 seconds. Animals were tested sequentially at 30 °C, 40 °C, and 50 °C, advancing to the next temperature only if no response occurred. No mice, either Sall3^cKO^ or control littermates, responded at 30 °C or 40 °C, and all mice responded at 50 °C.

#### CatWalk

Gait analysis was performed using the CatWalk XT system (Noldus Information Technology), a video-based platform that quantifies paw placement and locomotor dynamics. Mice traversed an illuminated glass runway while a high-speed camera positioned beneath the walkway captured paw contacts. The CatWalk software (v10) automatically identified paw prints and extracted a range of spatiotemporal gait parameters, including stride length, swing and stance duration, base of support, interlimb coordination, paw intensity/pressure, and regularity index. Automated paw assignments were reviewed for accuracy, and corrections were made when necessary. Each mouse completed multiple runs across the runway until three compliant runs (consistent speed with no pauses or grooming) were obtained for analysis. For each mouse, values from the three compliant runs were averaged to generate a single measurement per parameter.

#### Rotarod

Motor coordination and balance were assessed using an accelerating rotarod (Med Associates, model# ENV-575M). On the training day, mice received three trials on a rod rotating at a constant 4 rpm for 1 minute or until they fell. On the testing day, mice completed three trials in which the rod accelerated from 4 to 40 rpm. A 15-minute inter-trial interval was provided between all trials. For each mouse, the latency to fall from the three testing-day trials was averaged to generate a single performance value.

#### Pellet Reaching

A single-pellet forelimb reaching task was performed as previously described.^69^ To motivate reaching behavior, mice were placed on a restricted feeding schedule beginning prior to training and maintained throughout the experiment such that body weight remained at 85-90% of baseline. The training apparatus consisted of a Plexiglass box with a vertical slit opening, as described previously.^70^ A platform positioned in front of the opening held a small food pellet (Bio-Serv, cat# F0071) for the mouse to retrieve.

During the week prior to training, mice were habituated to the apparatus with food pellets provided inside the chamber. Paw preference was then determined by placing a small pile of pellets on the platform and allowing mice to perform 10-15 reaching attempts. The preferred paw was defined as the paw used most frequently or when the same paw was used for five consecutive reaches.

Training consisted of 10 daily sessions, each lasting until either 30 reaching attempts were completed or 20 minutes had elapsed. Reaches were manually recorded and classified as successful when the mouse used its preferred paw to grasp the pellet and bring it directly to its mouth. Success rate was calculated as the percentage of successful reaches out of total reaching attempts. Mice that failed to perform reaching attempts during training were excluded from analysis.

### MoSeq

Data acquisition was performed as previously described.^71^ The enclosure (diameter: 18”, wall height: 15”) was painted black and sanded to avoid image artifacts and reflections. Mice were placed in the center of the enclosures and allowed to freely explore for 20 minutes under red-light illumination in an otherwise dark environment. Each mouse was tracked in 3D with Azure Kinect camera (Microsoft) attached above the recording arena at a working distance of roughly 23.5 inches. Raw data from the camera was transmitted to an acquisition computer (Thermaltake Glacier 360 Liquid-Cooled PC with an AMD Ryzen 5 5600X, GeForce RTX 3060, 16GB 300 Mhz DDR4 ToughRAM RGB Memory and 1TB NvMe M.2) via USB 3.0 cables, and depth frames were retrieved at 30 Hz via the azure-acquire Python package (v2023.1.0)^71^ and saved to disk via FFV1-encoded videos.

Data preprocessing, extraction, and modeling were performed as previously described^71–74^ on local machines. Raw depth data was processed to extract the mouse’s position, orientation, and 3D profile. Each recording was first analyzed to compute a median background image of the empty area, which was then subtracted from each frame to isolate the mouse. The resulting 3D images of the mouse were cropped and aligned using a pre-trained classifier to ensure consistent orientation, with the mouse’s head always facing right. A Principle Component Analysis (PCA) was used to reduce the dimensionality of the extracted depth video, and covariance between the resulting principle components (PCs) was removed using Cholesky whitening. Ten PCs were retained for modeling as they explain 90% of the variance. To evaluate the optimal kappa hyperparameter, the whitened PCs were fit to 10 AR-HMMs through Gibbs resampling for 100 iterations, with kappa values linearly spaced between 1,000 and 100,000. Models were compared based on their median log-likelihood, and the model with the optimal kappa was fit for 1,000 iterations. The set of syllables identified by the final AR-HMM was filtered down to 42 syllables, which captured 99% of the variance.

Syllables were characterized by two objective kinematic parameters extracted directly from the MoSeq pipeline: mean displacement (px) and mean velocity (mm/frame). Each syllable was plotted in this two-dimensional kinematic space and partitioned into four groups based on whether its mean displacement and mean velocity fell above or below the median of each parameter across all syllables. This yielded four kinematically-defined syllable groups: high-displacement/high-velocity, high-displacement/low-velocity, low-displacement/high-velocity, low-displacement/low-velocity. Convex hulls were computed for each group to visualize the boundaries of each cluster, and centroids were calculated as the mean displacement and mean velocity across all syllables within each group. Syllable usage was quantified from the state sequence output of the AR-HMM. For each recording, the usage of each syllable was computed as the proportion of frames assigned to that syllable, and syllable usages were then averaged across animals within each experimental group.

#### Tapered Beam

Skilled motor coordination was assessed using a 100-cm tapered beam that narrowed from 3.5 cm at the starting end to 0.5 cm at the opposite end.^75,76^ The beam was elevated above the tabletop with the wide end positioned 32 cm above the surface and the narrow end positioned 7 cm above the surface, resulting in a gentle incline (16.2° at the narrow end; 73.8° at the wide end). Centimeter markings along the beam allowed for positional scoring. An enclosed dark chamber with a 2 x 2 inch opening was placed at the narrow end to provide a consistent endpoint for traversal.

Mice were acclimated to handling and the testing environment for two days (approximately 1 hour per day). During subsequent habituation and testing, mice traversed the beam three times per session. All runs were recorded using a video camera (EOS Rebel T5i, Cannon), and videos were scored for traversal time as well as the number and spatial distribution of foot faults. For each mouse, values from three successful trials were averaged to generate a single performance metric.

#### Swimming

Swimming behavior was assessed in a rectangular glass tank (48” x 4” x 4”) filled to approximately two-thirds of its height with water maintained at 23-25 °C. Mice were gently lowered by the tail into one end of the tank and allowed to swim toward a visible escape platform at the opposite end. Animals were acclimated to the testing room for at least 60 minutes before testing and were assessed under normal room lighting. Each mouse performed multiple swim trials, which were recorded using an iPhone 13 in slow-motion mode. Between trials, mice recovered in a clean cage positioned partially on a 37 °C heating pad, allowing voluntary thermoregulation. After the final trial, mice were dried as needed and returned to their home cage.

For kinematic analysis, one representative video per mouse was converted to AVI format and imported into ImageJ (v1.54p). Frames were treated as a z-stack, and 2D coordinates were collected using the Cell Counter plugin for the following landmarks: base of tail, tip of tail, left and right forelimbs, and left and right hindlimbs. Base and tip of tail were annotated in every frame. Forelimbs were marked at their most ventral point when used for propulsion. Hindlimbs were annotated across each stroke, with one frame intentionally skipped between strokes to facilitate identification of continuous swing segments.

Annotations were processed in Python using custom scripts. The analysis produced four classes of metrics:

1. *Forelimb reach heatmaps*. Frame indices were normalized to swim duration (0–1) and divided into 20 equal bins. For each bin, the fraction of frames in which either forelimb was engaged was computed. These per-mouse fractions were visualized as heatmaps, with rows ordered by each mouse’s total number of either-forelimb engagement events.
2. *Tail position*. The vertical position of the tail tip was expressed relative to the nose in the vertical direction, where 0 indicates the tail tip is level with the nose and positive values indicate the tail tip is above the nose. Tail-tip values were normalized in time (0–1) and binned into 20 intervals. For each mouse and bin, the maximum tail height within each bin was calculated and displayed as a heatmap ordered by each mouse’s mean maximum tail height across bins.
3. *Foot displacement*. Horizontal displacement of the left paw relative to the base of the tail was computed for each swing (positive when the foot was ahead of the tail base, negative when behind). Each swing was phase-normalized to 0–1 and resampled to 101 points. For each mouse, displacement traces were averaged across swings, and the genotype-level means with 95% confidence intervals were plotted against swing phase.
4. *Left-right alteration.* For each swing, the onset times of left and right hindlimb swings were compared to generate a fractional phase offset (0–1), normalized by the stride period. Values near 0.5 indicate near-perfect alteration. Each mouse’s distribution of offsets was binned into 10% intervals and visualized as heatmaps, with rows ordered by each mouse’s mean phase offset.

### Electrophysiology

The procedures for *ex vivo* sacrocaudal spinal cord preparation have been previously described.^77,78^ Animals were deeply anesthetized with urethane (≥0.2 g/100 g, i.p.) and supplied with carbogen (95% O_2_ / 5% CO_2_) through a face mask. With the animal in sternal recumbency, a dorsal laminectomy was performed to expose the lower half of the spinal cord. The cord was transected at the lumbosacral enlargement (L5–L6), and the entire sacrocaudal segment (S1–Co2) with attached roots was transferred to a dissection dish containing artificial cerebrospinal fluid (ACSF) continuously bubbled with carbogen at room temperature (∼21 °C). The ACSF contained (in mM): 128 NaCl, 3 KCl, 1.5 MgSO_4_, 1 NaH_2_PO_4_, 2.5 CaCl_2_, 22 NaHCO_3_, and 12 glucose (osmolarity ∼295 mOsm; pH 7.35–7.4 when aerated). Dorsal and ventral roots were separated and identified before the cord was transferred to a custom perfusion chamber.

In the chamber, the cord was positioned dorsal side up. On both sides of the cord, the S3 and S4 dorsal roots (DRs) were placed on bipolar wire electrodes connected to separate channels of an S88 Grass stimulator (A-M Systems) for conditioning and test stimulation, respectively. The S2 DR was attached to a suction electrode for recording dorsal root potentials (DRPs), reflecting primary afferent depolarization (PAD). A chlorinated silver wire wrapped around the electrode tip and immersed in ACSF served as the reference electrode. The S4 ventral root (VR) was mounted on bipolar wire electrodes connected to a differential amplifier to record motor output. All roots positioned on wire electrodes were covered with petroleum jelly to prevent drying. Preparations were allowed to recover in the perfusion chamber for approximately 1 hour before recordings began.

#### Electrophysiological Recordings

1. *Sensory-Motor Reflex*. The stimulation threshold (T) of the S4 DR was defined as the minimum current (0.1 ms pulse) that evoked the smallest detectable response in the S4 VR. Monosynaptic reflex curves were generated using stimulus intensities of 1, 1.25, 1.5, 2, 5, and 10×T delivered to the S4 DR. Stimulation was applied at 15 s inter-pulse intervals (0.067 Hz) to avoid adaptation^79,80^ and each stimulus intensity was repeated 5 times. The S4 VR and S2 DR were connected to differential amplifiers with a gain of 1000× to record ventral root compound action potentials (CAPs) and dorsal root potentials (DRPs), respectively. VR signals were recorded in AC mode with a 300 Hz-3 kHz bandwidth, whereas DRPs were recorded in DC mode and filtered at 0.1 Hz-3 kHz. Signals were digitized using a Micro3-1401 interface (CED, UK) at 10-20 kHz and acquired in Spike2® (Cambridge Electronic Design, v10.18) for offline analysis. The monosynaptic S4 VR response was quantified as the peak-to-peak amplitude of the earliest CAP component. For each stimulus intensity, responses from typically five trials were averaged. PAD amplitude was measured as the peak of the average DRP recorded from the S2 DR. Trials containing dorsal root reflex (DRR) activity were excluded or low-pass filtered to prevent contamination of PAD measurements.
2. *Conditioned Monosynaptic Reflex*. The monosynaptic S4 ventral root response to S4 DR stimulation (test) was conditioned by prior stimulation of the S3 DR. The test stimulus was delivered to S4 at 2×T. The conditioning stimulus was delivered to S3 at an intensity equal to 2×T for evoking PAD in the S2 dorsal root, while ensuring it did not evoke a motor response in S4, as excessive afferent activation can recruit neighboring segments and produce motor discharge leading to post-activation depression of the test response.^45^ Conditioning and test stimuli were separated by inter-stimulus intervals (ISIs) of 10, 20, 30, 50, 100, 200, 1000, and 2000 ms, with each ISI repeated five times. Conditioned monosynaptic CAP amplitudes were quantified as peak-to-peak values and, within each pharmacological condition, normalized to the average unconditioned response evoked by S3 DR stimulation alone.
3. *Rate-Dependent Depression (RDD)*. RDD was assessed by delivering trains of five consecutive pulses to the S4 dorsal root at stimulation frequencies of 0.67, 1, 5, 10, 25, 50, and 100 Hz, using stimulation intensities of 1.25, 1.5, 2, 5, and 10×T. For each frequency and stimulation intensity condition, the percent decrease in CAP amplitude relative to the first pulse was calculated for pulses 2–5 and then averaged to obtain a single RDD value.

Drug Application. All recording paradigms were first obtained in ACSF and then repeated following application of the GABA_B_ receptor blocker (CGP 55485, 2 µM; Tocris, cat# 1248) and the GABA_A_ receptor blocker (gabazine, 20 µM; Tocris, cat# 1262), with a 30 min washout period between drug applications.

### Quantification and statistical analysis

#### Derived measures

Cell-type percentages were calculated relative to the indicated parent population. Sall3 expression was normalized within each anatomical region to the mean control value, which was set to 100%.

The percentage of vGlut1 terminals containing GAD65on signal was calculated as the number of vGlut1 surfaces associated with GAD65on signal divided by the total number of vGlut1 surfaces, multiplied by 100.

Mean fluorescence intensities were quantified within Imaris-defined surfaces for each section. To account for differences in baseline fluorescence intensity across experimental batches, least-squares mean predictions and residuals from the mixed-effects model described in Table S1 were saved and summed to generate batch-corrected log10 intensity values for each section. Corrected values were averaged across sections for each mouse and anatomical region. Group means ± SEM were plotted as a function of gene dose, with lines representing fitted trends and corresponding confidence intervals derived from the model.

For PAD and VRR stimulus-response curves, area under the curve (AUC) was calculated for each preparation by trapezoidal integration of response amplitude across the available stimulus intensities from 1 to 10×T. Missing measurements were omitted, and AUC was calculated only when measurements were available at four or more stimulus intensities.

For conditioned monosynaptic reflex experiments, the effect of the conditioning stimulus was expressed as the percentage change from the corresponding unconditioned test response: ΔMSR (%) = 100 × (conditioned MSR − unconditioned MSR)/unconditioned MSR. Positive AUC over the 0–100 ms interstimulus-interval window (AUC_0–100_) was calculated by trapezoidal integration, including only portions of the response above zero. When a line segment crossed zero, the crossing point was determined by linear interpolation, and only the positive portion of the segment was included.

For rate-dependent depression, each response was expressed as the percentage change from the first pulse: ΔMSR (%) = 100 × (Pn − P1)/P1. Values from pulses 2–5 were averaged to generate a single RDD value for each stimulation frequency and intensity. Drug-induced changes in RDD were calculated by subtracting the corresponding ACSF value from the value obtained following CGP or gabazine treatment.

For tapered-beam spatial analysis, the beam was divided into four equal-length regions, and foot faults within each region were averaged across trials for each mouse. A simple linear regression was fit to foot-fault number as a function of ordered beam region (1–4), and the resulting slope was used as the per-mouse measure of foot-fault distribution.

For swimming kinematics, maximum forward and reverse displacement were defined as the maximum and minimum values, respectively, of each mouse’s phase-normalized mean displacement trace. Swing amplitude was calculated as maximum forward displacement minus maximum reverse displacement.

#### Statistical analysis

Statistical analyses were performed using GraphPad Prism (v10.5.0) and JMP Student Edition (v18). The individual mouse was the experimental unit for anatomical and behavioral analyses, and the individual spinal cord preparation was the experimental unit for electrophysiological analyses. Data are presented as mean ± SEM unless otherwise indicated, with individual data points representing independent animals or preparations.

Two-group comparisons were performed using two-tailed paired or unpaired t tests, with Welch’s correction when unequal variances were present. Mann–Whitney tests were used for non-normally distributed data or data containing zero values that precluded an appropriate transformation. Comparisons involving multiple groups or factors were performed using one- or two-way ANOVA. Repeated-measures ANOVA or mixed-effects models were used when measurements were matched within animals or preparations. Survival distributions were compared using the log-rank (Mantel–Cox) test. Associations between continuous variables were assessed using simple linear regression, and regression slopes were compared using extra sum-of-squares F tests. Multiple-comparison procedures included Tukey’s, Šídák’s, Dunnett’s, Dunnett’s T3, and Holm–Šídák tests, as appropriate.

When necessary, data were log10-, log10(x + 1)-, or square-root-transformed to reduce right-skew or accommodate zero values. All tests were two-sided, and p < 0.05 was considered statistically significant. The statistical test, transformation, multiple-comparison procedure, exact p values, sample sizes, means, and standard deviations for each analysis are provided in Table S1.

## Supporting information

Table S2 - Statistics

Table S2 - Brain Sall3 Expression

Video S1 - Tapered Beam Crossing

Video S2 - Pellet-Reaching

Video S3 - Swim Behavior

## Acknowledgements

We are grateful to C. Hansen for assistance with cryosectioning and microscopy. We thank S. Pfaff and P. Osseward for insightful discussions and for sharing their expertise, and J.M.H. for helpful and critical comments on the manuscript. We thank the Stanford Behavioral and Functional Neuroscience Laboratory (N. Saw, supervisor) for technical support with the CatWalk and rotarod assays. We also thank A. Nippert, L. Donovan, and V. Tawfik for sharing the von Frey protocol and analysis guidance, and O. Zhou and A. Brunet for providing the tapered beam protocol. We thank O.H. for assistance with early bioinformatic analysis, R.H.S. for assistance with Python programming and designing the cell-counting software, and J.P.H. for statistical consultation. This work was supported by NIH K99 NS133476 (M.A.G), New Jersey Commission on Spinal Cord Research (M.A.G and V.E.A), NIH/NINDS R01 NS119268 (V.E.A), NIH/NINDS K01 NS116224 (V.E.A), California Institute for Regenerative Medicine TRAN1-12891 & DISC2-12169 (I.L.L), NIH R01 NS083998 (J.A.K), the Wu Tsai Neurosciences Institute (J.A.K), and the Stanford University Department of Neurosurgery (J.A.K).

## Author Contributions

J.L.S. and J.A.K. conceived the study. J.L.S. performed the single-cell analysis, synapse quantification, and behavioral experiments. R.P. and T.W-C. organized and curated the sequencing data to enable downstream analysis. Z.S. assisted with behavioral experiments. R.H.R. performed and analyzed the pellet reach motor assay. A.P. conducted MoSeq behavioral data acquisition and contributed to MoSeq analysis with analytical direction from V.K. and I.L.L. M.A.G. and V.E.A. provided Rorb mouse tissue. A.A.M. designed and conducted the electrophysiology experiments and analyzed the data with guidance from D.B. and C.J.H. J.P.G. provided statistical consultation. J.L.S. wrote the original draft, and J.L.S. and J.A.K. reviewed and edited the manuscript.

## Competing interests

Authors declare no competing interests

---

**Figure S1.**
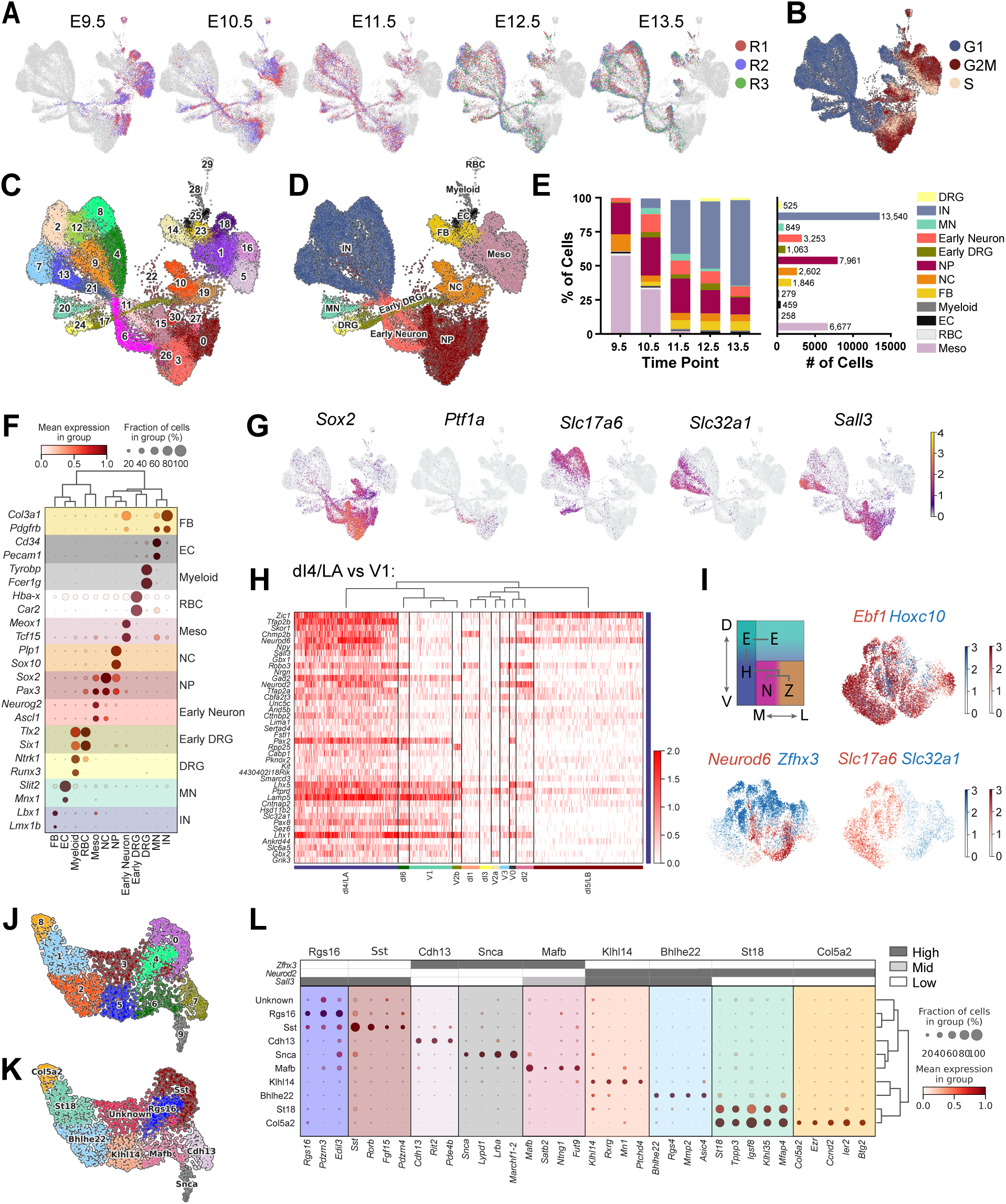
Single-cell transcriptomic profiling of developing spinal cord cells, related to Figure 1. **(A-B)** UMAP projections showing all cells from the Delile et al. developmental spinal cord dataset,^26^ colored by embryonic age across replicates R1-R3 (A) and by cell cycle status (B). Biological replicates show strong overlap, while cells segregate by developmental age, indicating minimal batch effects. Cell cycle analysis distinguishes proliferative neural progenitors from postmitotic neuronal populations, including a population of ‘early neurons’ that have exited the cell cycle but have not yet acquired canonical excitatory or inhibitory markers. **(C-D)** UMAP projections showing cluster assignments (C) and annotated cell type identities (D). **(E)** Proportion of all cells at each time point (left) and total cell number (right) for each annotated cell cluster. **(F)** Dot plot of canonical transcription factor expression across annotated cell types. **(G)** Expression of *Sox2* (NP), *Ptf1a* (early neurons), *Slc32a1* (inhibitory vGAT interneurons), and *Slc17a6* (excitatory vGlut2 interneurons) in UMAP space. *Sall3* expression is enriched in inhibitory interneurons and is also detected in early neurons and neural progenitors. **(H)** Heatmap of genes enriched in dI4/LA relative to V1, with dI4/LA-enriched genes displayed across annotated interneuron classes at single-cell resolution. The top 40 filtered genes with positive log fold change are shown. **(I)** UMAPs showing hierarchical transcription factor organization of interneurons, based on the E-H (*Ebf1*, *Hoxc10*) and N-Z (*Neurod2*, *Zfhx3*) classification framework described by Osseward et al.^33^ Top left: Schematic illustrating relationships between E-H and N-Z marker expression along the dorsal-ventral (D-V) and medial-lateral (M-L) axes. Not shown, H-, N-, and Z-group populations are further subdivided by neurotransmitter identity into excitatory (*Slc17a6*) and inhibitory (*Slc32a1*) subtypes. Expression of E-H (top right), N-Z (bottom left), and excitatory-inhibitory (bottom right) markers is shown. Consistent with prior reports, E-H subtypes are not fully resolved at early developmental stages, whereas N-Z segregation emerges along the y-axis of the UMAP and neurotransmitter identity separates along the x-axis. **(J-K)** dI4/LA interneurons from E12.5 and E13.5 samples were reclustered for further analysis, with resulting cluster structure displayed in UMAP space for all clusters (J) and annotated clusters (K). **(L)** Dot plot of annotated dI4/LA interneurons showing marker expression across subpopulations. The Klhl14, Bhlhe22, St18, and Col5a2 clusters correspond to N-group neurons (*Neurod2*^+^), which are typically later-born and positioned more dorsomedially. Cdh13, Snca, and Mafb mark Z-group neurons (*Zfhx3*^+^), a population generally associated with earlier neurogenesis and more lateral settling. Rgs16 and Sst express *Sall3* but show low N- or Z-group marker expression, suggesting they may represent E-group neurons in the superficial dorsal horn. Within N-group neurons, *Sall3* expression is restricted to the Bhlhe22 and Klhl14 clusters. Given prior studies linking *Klhl14* to GABApre interneurons,23 this cluster is consistent with the dI4/LA subset contributing to GABApre circuitry. Abbreviations: IN, interneurons; MN, motor neurons; DRG, dorsal root ganglion; NP, neural progenitor; NC, neural crest; Meso, mesoderm; FB, fibroblast; EC, endothelial cell; RBC, red blood cell.

**Figure S2.**
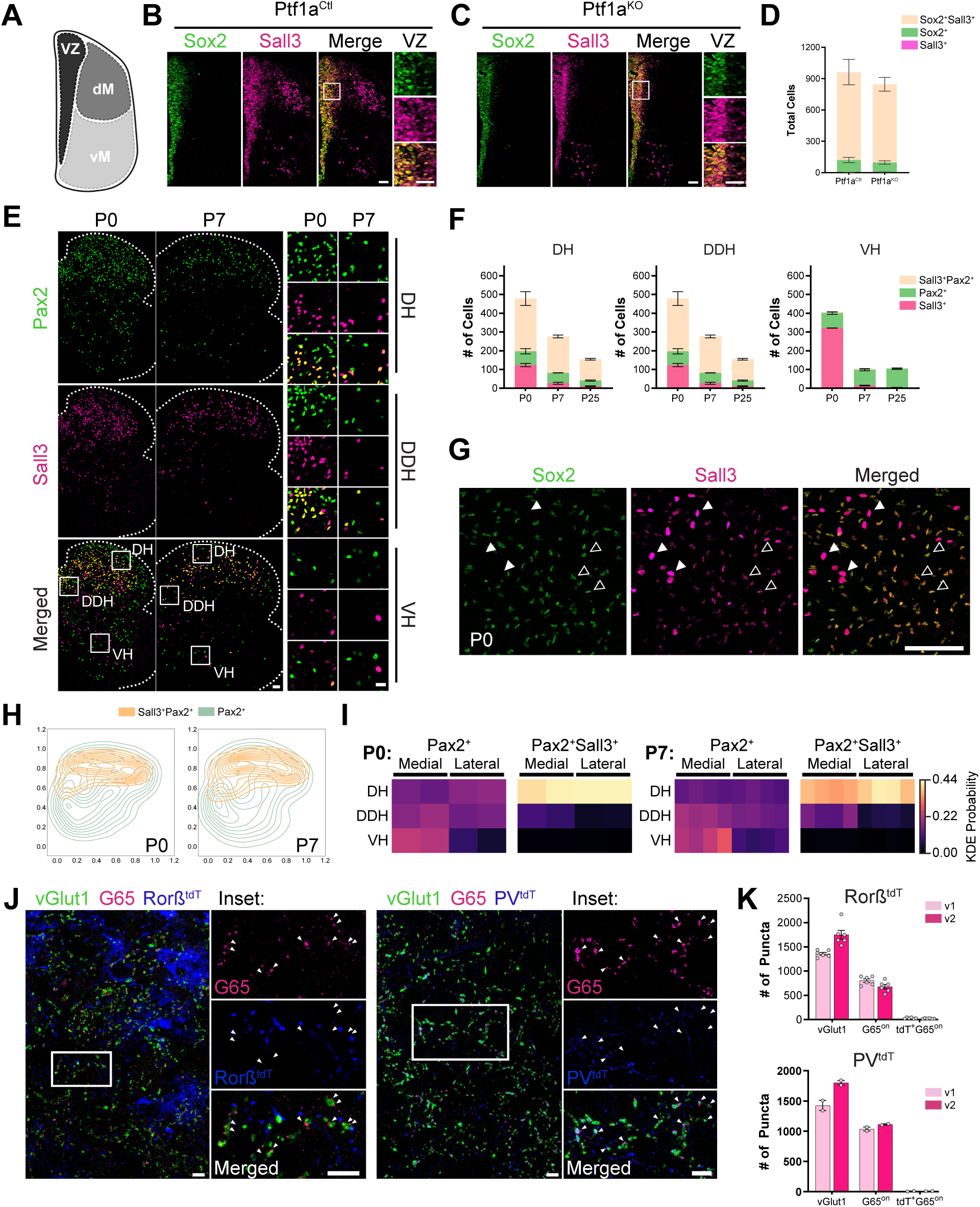
Developmental distribution and cellular identity of Sall3^+^ cells in the postnatal spinal cord, related to Figure 2. **(A)** Schematic of the embryonic spinal cord. **(B-C)** Immunostaining for Sox2 (green) and Sall3 (magenta) in representative Ptf1a^Ctl^ (B) and Ptf1a^KO^ (C) E12.5 embryonic spinal cord sections. Insets show higher-magnification views of the VZ. Sall3 is expressed in most Sox2^+^ neural progenitors, consistent with single-cell sequencing analysis (see Figure S1G). **(D)** Quantification of Sox2 and Sall3 co-expression in the VZ of Ptf1a^Ctl^ and Ptf1a^KO^ embryos. **(E)** Immunostaining for Pax2 (green) and Sall3 (magenta) in representative P0 and P7 spinal cord hemisections. Insets show higher-magnification views of the DH, DDH, and VH at each time point. **(F)** Stacked bar plots showing total cell counts per section in DH (left), DDH (middle), and VH (right). Cell classes include: Sall3^+^Pax2^+^ (peach), Pax2^+^ (green), and Sall3^+^ (magenta). **(G)** Immunostaining for Sox2 (green) and Sall3 (magenta) in representative P0 spinal cord sections. Following the gliogenic switch, Sox2^+^ progenitors give rise to astrocytes that retain Sox2 expression. Large, bright Sall3^+^Sox2^-^ cells correspond to neurons (closed arrowheads), whereas smaller, dimmer Sall3^+^Sox2^+^ cells correspond to astrocytes (open arrowheads). Dim Sox2^+^Sall3^+^ astrocytes are detected at early postnatal stages and decrease with age (see pink bar in Figure S2F). **(H-I)** Spatial distribution of Sall3^+^Pax2^+^ (orange) and Pax2^+^ (green) interneurons at P0 and P7. Contour plots (H) and corresponding heatmaps (I) show kernel density estimation (KDE) probability across DH, DDH, and VH, as in Figure 2. **(J)** Immunostaining for vGlut1 (green), G65 (magenta), and tdTomato (blue) in representative P25 ventral horn (v1) spinal cord sections from Rorb^tdT/+^ (left) and PV^tdT^ (right) mice. Insets show higher-magnification views of the indicated regions. Arrowheads highlight absence of tdT expression in G65^+^ GABApre boutons. **(K)** Quantification of vGlut1^+^ sensory terminals, G65^on^ boutons, and tdT^+^G65^on^ boutons in Rorb^tdT/+^ (top) and PV^tdT^ (bottom) mice. Scale bars: (B and C) 50 µm (overview and inset), (E) 50 µm (overview), 25 µm (insets), (G) 100 µm, (J) 10 µm (overview and inset). Graphs show mean ± SEM, with dots representing individual mice. Exact sample sizes (N), means, and standard deviations are provided in Table S1. Abbreviations: VZ, ventricular zone; dM, dorsal mantle zone; vM, ventral mantle zone; DH, dorsal horn; DDH, deep dorsal horn; VH, ventral horn.

**Figure S3.**
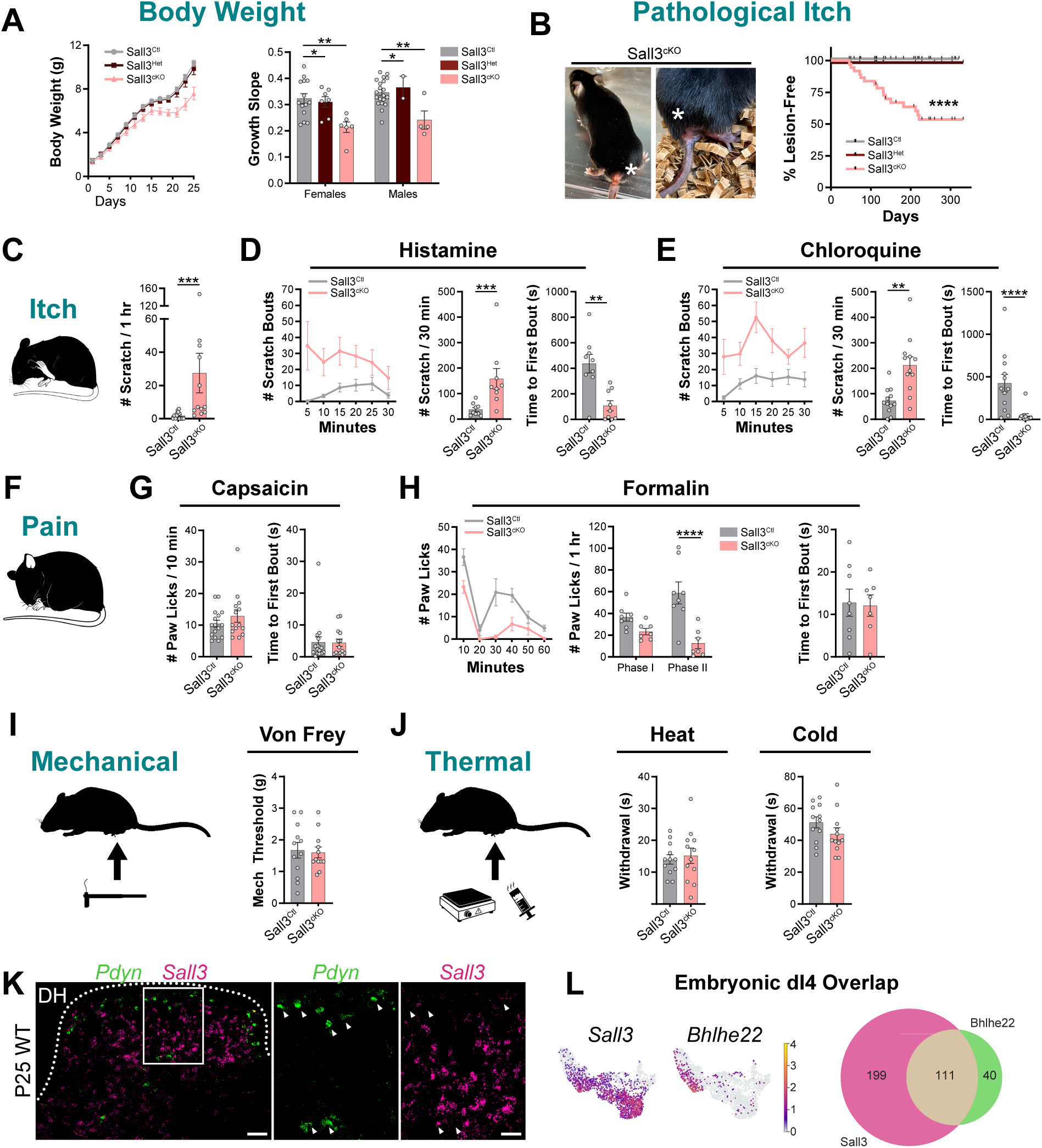
Sensory phenotype of Sall3 conditional mutants, related to Figure 2. **(A)** Body weight measurements from P1-P25 in Sall3^Ctl^, Sall3^Het^, and Sall3^cKO^ mice (left). Growth slope analysis separated by sex (right). **(B)** Representative images of Sall3^cKO^ mice exhibiting lesions near the base of the tail (left). Kaplan-Meier plot showing the percentage of lesion-free mice across genotypes (right). **(C)** Illustration of scratching behavior during itch responses and quantification of spontaneous scratching behavior during a 60 min recording session. Total number of scratch bouts is shown for Sall3^Ctl^ and Sall3^cKO^ mice. **(D)** Quantification of histamine-induced itch. Scratch bouts binned in 5 min intervals across a 30 min session (left), total number of scratch bouts (middle), and latency to first scratch bout (right). **(E)** Quantification of chloroquine-induced itch. Plots displayed as in (D). **(F)** Illustration of paw-lick behavior following application of noxious stimuli. Lick behavior was recorded during 10 min sessions following capsaicin injection or 60 min sessions following formalin injection. **(G)** Quantification of capsaicin-induced nociception. Total number of paw licks (left) and latency to first lick (right). **(H)** Quantification of formalin-induced nociception. Plots displayed as in (D), with the total number of paw licks separated as phase I (first 10 min) and phase II (remaining 50 min). **(I)** Illustration of the von Frey mechanical sensitivity assay and quantification of the withdrawal threshold. **(J)** Illustration of thermal sensitivity assays using hot plate (heat) and dry ice (cold) paradigms. Latency to foot withdrawal on the hot plate (left) or after exposure to dry ice administered under a glass pane (right). **(K)** *In situ* hybridization for *Prodynorphin* (*Pdyn*; green) and *Sall3* (magenta) in a representative P25 wild-type dorsal horn (DH). Inset shows a higher-magnification view of the indicated region. Arrowheads highlight absence of *Sall3* expression in *Pdyn*^+^ interneurons. **(L)** UMAP projections of *Sall3* and *Bhlhe22* expression within embryonic dI4/LA interneuron subpopulations (left). Venn diagram showing overlap between *Sall3*- and *Bhlhe22*-expressing cells within the *Bhlhe22* cluster (right). Scale bars: (K) 50 µm (overview), 25 µm (insets). Graphs show mean ± SEM, with dots representing individual mice. Statistical significance is indicated as ** *p* < 0.01, *** *p* < 0.001, **** *p* < 0.0001. Exact *p* values, sample sizes (N), means, and standard deviations are provided in Table S1.

**Figure S4.**
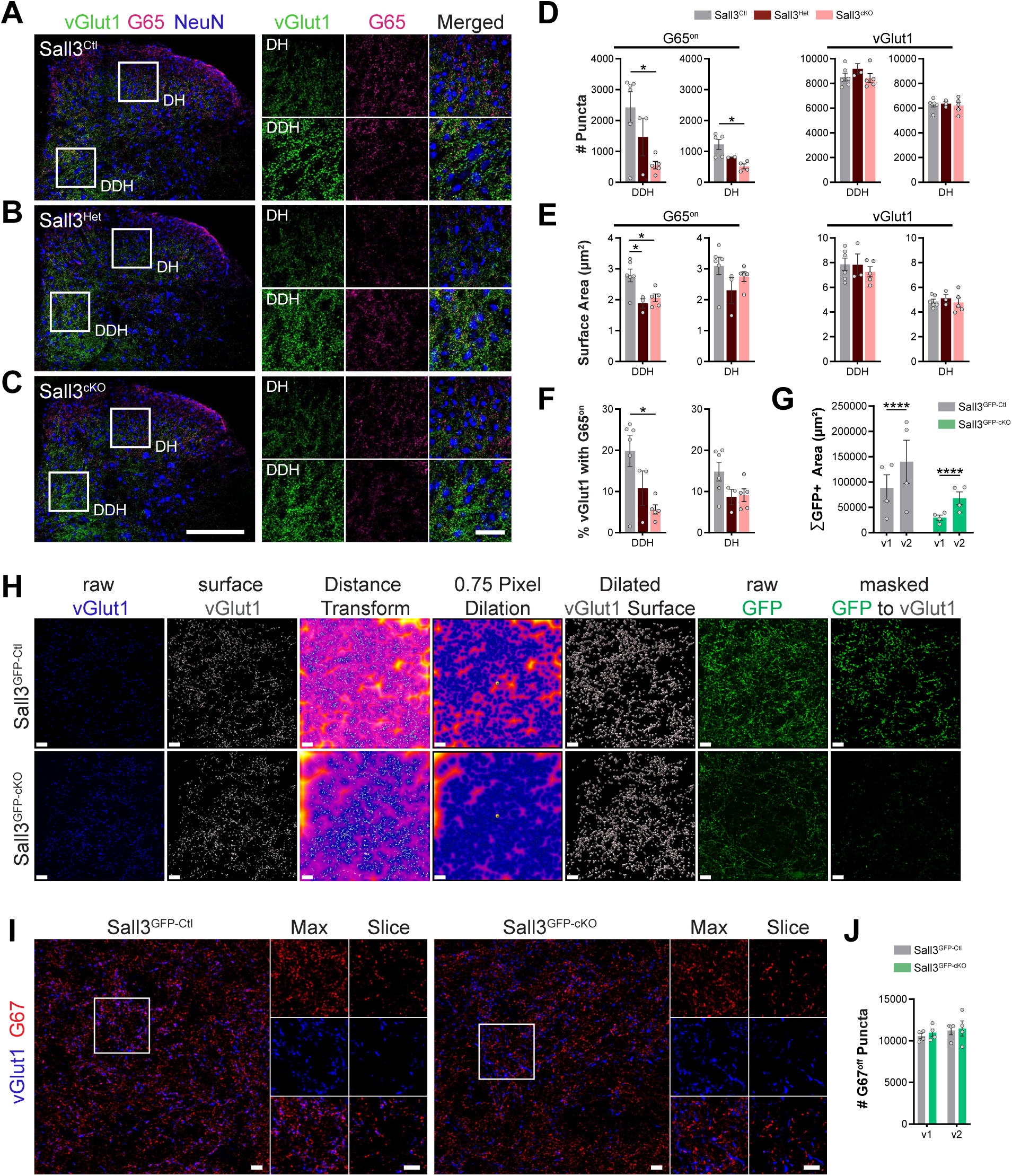
Sall3-dependent loss of GABApre boutons extends to the dorsal spinal cord, related to Figure 3. **(A-C)** Immunostaining for vGlut1 (green, sensory afferent terminals), G65 (GAD65; magenta, presynaptic boutons), and NeuN (blue, neurons) in more dorsal regions of representative Sall3^Ctl^ (A), Sall3^Het^ (B), and Sall3^cKO^ (C) spinal cord sections. Insets show higher-magnification views of DH and DDH. **(D-E)** Quantification of synapse reconstructions from DH and DDH regions across all genotypes. The number of puncta per section (D) and mean surface area (E) are shown for G65^on^ and vGlut1 puncta. **(F)** Percentage of vGlut1 surfaces contacting at least one associated G65^on^. **(G)** Total surface area of GFP^+^ projections in the v1 and v2 regions of Sall3^GFP-Ctl^ and Sall3^GFP-cKO^ mice. **(H)** Workflow illustrating the synapse reconstruction and masking strategy for Gad2^GFP-Ctl^ (top) and Gad2^GFP-cKO^ (bottom) mice. Panels show raw vGlut1 signal (blue), vGlut1 surface reconstruction (gray), distance transformation of the vGlut1 surface, dilation of the distance transformation, generation of a dilated vGlut1 surface, raw GFP signal (green), and GFP signal masked to the dilated vGlut1 surface. **(I)** Immunostaining for G67 (GAD67; red, GABApost boutons) and vGlut1 (blue, Ia afferent terminals) in the v1 region of the ventral horn in representative Sall3^GFP-Ctl^ (left) and Sall3^GFP-cKO^ (right) spinal cord sections. Insets show higher-magnification views of both maximum intensity projections (Max) and single optical sections (Slice). **(J)** Quantification of G67^off^ (GABApost) synapses in v1 and v2 regions from Gad2^GFP-Ctl^ and Gad2^GFP-cKO^ mice based on synapse reconstructions. Scale bars: (A-C) 100 µm (overview), 50 µm (insets); (G) 20 µm; (H) 10 µm (overview and insets). Graphs show mean ± SEM, with dots representing individual mice. Statistical significance is indicated as ** *p* < 0.01. Exact *p* values, sample sizes (N), means, and standard deviations are provided in Table S1.

**Figure S5.**
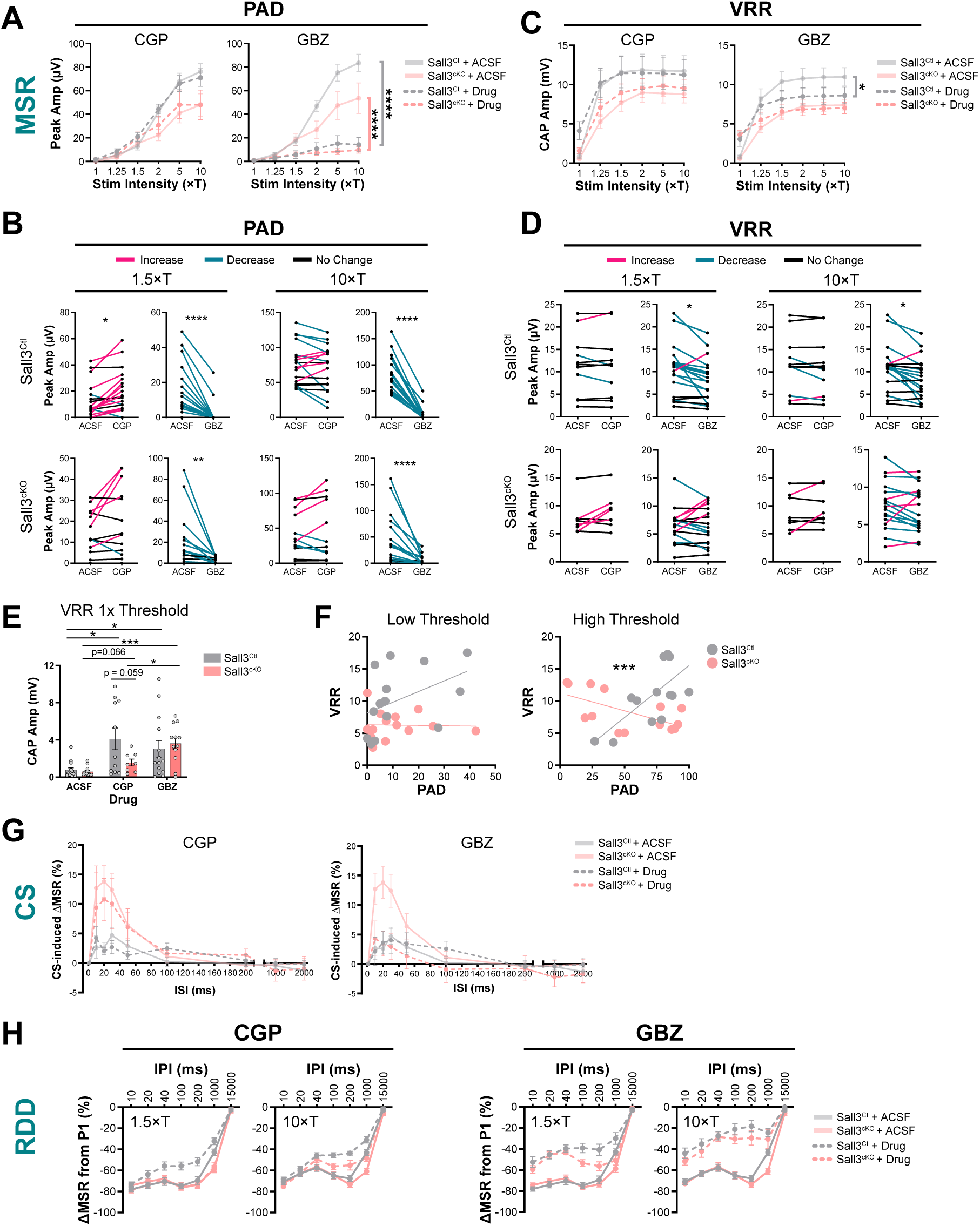
Expanded analyses of GABA_A_-mediated PAD and GABA_B_-mediated presynaptic inhibition in Sall3^cKO^ mice, related to Figure 5. **(A)** Quantification of PAD peak amplitude across stimulus intensities in Sall3^Ctl^ and Sall3^cKO^ mice. ACSF baseline responses (replotted from Figure 5C for comparison) are shown alongside the effects of CGP (left) and GBZ (right). **(B)** Paired comparisons of PAD peak amplitude under ACSF baseline conditions and following CGP or GBZ application at 1.5×T (left) and 10×T (right) stimulation intensities. Sall3^Ctl^ mice are shown in the top row and Sall3^cKO^ mice in the bottom row. Pairs are colored magenta for an increase, blue for a decrease, and black for no change. **(C)** Quantification of VRR CAP amplitude across stimulus intensities in Sall3^Ctl^ and Sall3^cKO^ mice. ACSF baseline responses (replotted from Figure 5D for comparison) are shown alongside the effects of CGP (left) and GBZ (right). **(D)** Paired comparisons of VRR CAP amplitudes under ACSF baseline conditions and following CGP or GBZ application at 1.5×T (left) and 10×T (right) stimulation intensities. Sall3^Ctl^ mice are shown in the top row and Sall3^cKO^ mice in the bottom row. **(E)** Quantification of VRR responses at 1×T, where only a small number of Ia proprioceptors are recruited. CAP amplitude increases under CGP and GBZ blockade in Sall3^Ctl^ mice, whereas in Sall3^cKO^ mice the response increases with GBZ and shows reduced sensitivity to CGP. **(F)** Correlation between PAD and VRR responses in Sall3^Ctl^ and Sall3^cKO^ mice at low (1.25-1.5×T; left) and high (5-10×T; right) stimulation intensities. **(G)** Quantification of test-evoked CAP amplitudes across ISIs under CGP (left) and GBZ (right) conditions. Test responses are expressed as the percent change from the corresponding unconditioned response. ACSF baseline curves (replotted from Figure 5G for comparison) are shown alongside drug-treated responses. **(H)** Quantification of rate-dependent depression (RDD) showing monosynaptic reflex responses across ISIs under CGP (left) and GBZ (right) conditions. Responses are expressed as the percent change from the first pulse (P1). The 1.5xT and 10xT stimulation intensities are shown in separate panels. Graphs show mean ± SEM, with dots representing individual hemicord recordings analyzed as independent samples. Statistical significance is indicated as * *p* < 0.05, *** *p* < 0.001. Exact *p* values, sample sizes (N), means, and standard deviations are provided in Table S1.

**Figure S6.**
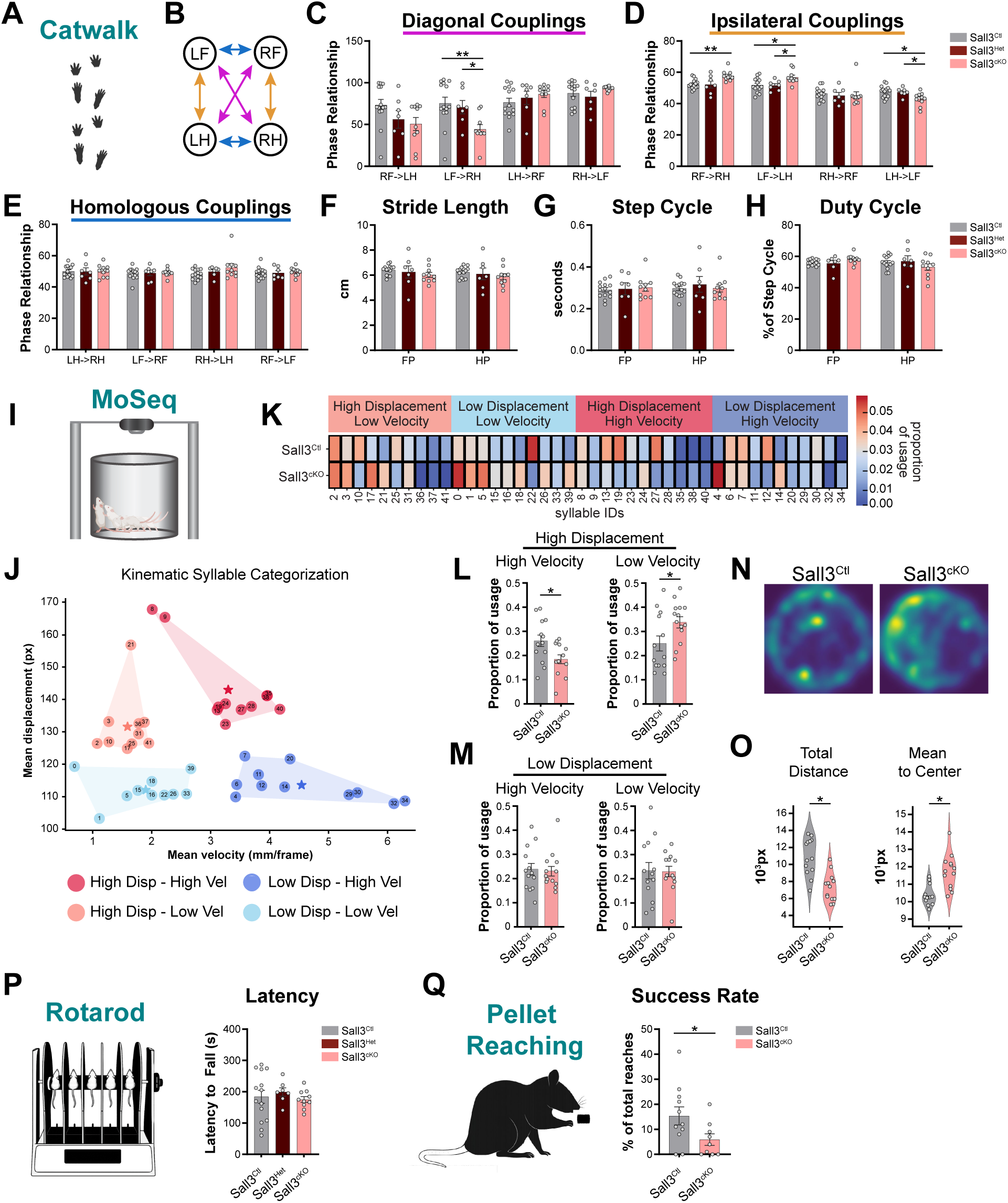
Expanded motor and coordination analyses, related to Figure 6. **(A)** Illustration of the CatWalk XT gait analysis system. **(B-E)** Quantification of CatWalk coupling relationships. Schematic illustrating diagonal (magenta), ipsilateral (yellow), and homologous (blue) couplings (B). Diagonal (C), ipsilateral (D), and homologous (E) interlimb phase relationships. Phase relationship values (0-100) represent the timing of the second paw contact relative to the first within the normalized stride cycle. **(F-H)** Quantification of CatWalk gait parameters. Stride length, defined as the distance traveled by a paw between two successive contacts of the same paw (F), step cycle, defined as the time between two successive contacts of the same paw (G), and duty cycle, defined as the percentage of the step cycle during which the paw is in contact with the ground (H). **(I)** Illustration of the Motion Sequencing (MoSeq) behavior setup. Mice freely explored an open area while behavior was recorded with an overhead depth camera and segmented into discrete behavioral syllables. **(J-M)** Behavioral syllables were categorized by mean displacement and mean velocity into high displacement/high velocity, high displacement/low velocity, low displacement/high velocity, and low displacement/low velocity groups (J). Heatmap showing usage of categorized behavioral syllables in Sall3^Ctl^ and Sall3^cKO^ mice (K). Quantification of syllable usage across high displacement (L) and low displacement (M) behavioral groups, separated by high and low velocity. **(N-O)** MoSeq occupancy plots showing spatial position during open-field exploration (N) and quantification of open-field locomotion showing total distance traveled (O, left) and mean distance to the center of the behavioral chamber (O, right). **(P)** Illustration of the rotarod apparatus and quantification of performance. Latency to fall during acceleration (4-40 rpm) was averaged across three trials. **(Q)** Illustration of the pellet-reaching task and quantification of performance. The percentage of successful reaches was averaged across three post-training test days. Graphs show mean ± SEM, with dots representing individual mice. Statistical significance is indicated as * *p* < 0.05, ** *p* < 0.01. Exact *p* values, sample sizes (N), means, and standard deviations are provided in Table S1. Abbreviations: RF, right front; LF, left front; RH, right hind; LH, left hind.

## Notes

### Competing Interest Statement

The authors have declared no competing interest.

